# Comparative landscape of genetic dependencies in human and chimpanzee stem cells

**DOI:** 10.1101/2023.03.19.533346

**Authors:** Richard She, Tyler Fair, Nathan K. Schaefer, Reuben A. Saunders, Bryan J. Pavlovic, Jonathan S. Weissman, Alex A. Pollen

## Abstract

Comparative studies of great apes provide a window into our evolutionary past, but the extent and identity of cellular differences that emerged during hominin evolution remain largely unexplored. We established a comparative loss-of-function approach to evaluate whether changes in human cells alter requirements for essential genes. By performing genome-wide CRISPR interference screens in human and chimpanzee pluripotent stem cells, we identified 75 genes with species-specific effects on cellular proliferation. These genes comprised coherent processes, including cell cycle progression and lysosomal signaling, which we determined to be human-derived by comparison with orangutan cells. Human-specific robustness to *CDK2* and *CCNE1* depletion persisted in neural progenitor cells, providing support for the G1-phase length hypothesis as a potential evolutionary mechanism in human brain expansion. Our findings demonstrate that evolutionary changes in human cells can reshape the landscape of essential genes and establish a platform for systematically uncovering latent cellular and molecular differences between species.

## Introduction

Comparative studies of humans and chimpanzees, our closest extant relatives, have long sought to define the evolutionary origins of unique human features. Within seven million years, humans evolved numerous specializations, from bipedalism to the threefold expansion of the cerebral cortex (Muller et al., 2017; Varki and Altheide, 2005). Many of these novel human traits emerge from changes in cell behavior during development. These changes in cell behavior may in turn create new requirements for existing genes and pathways that mediate evolutionary changes. However, we currently lack a framework for systematically identifying which molecular pathways play divergent roles in conserved developmental cell types.

Current approaches to studying the molecular basis of human evolution include reconstructing candidate mutations at specific loci in model organisms, but only a handful of mutations in non-coding regulatory regions and coding genes have been examined in detail. Among conserved non-coding elements with unexpected changes in the human lineage, specific loci have been linked to gene expression changes in distal limbs (Dutrow et al., 2022) increased sweat gland number (Aldea et al., 2021), and increased neural proliferation (Boyd et al., 2015). Among coding changes, two human-specific coding mutations in *FOXP2* have been proposed to contribute to human language capabilities based on functional studies in mouse models and human genetics (Enard et al., 2002; Lai et al., 2001), and three modern human-specific mutations in *KIF18A* and *KNL1* prolong metaphase and reduce segregation errors in neural progenitor cells (Mora-Bermúdez et al., 2022). In addition, recent duplications and subsequent modifications of *ARHGAP11B* and *NOTCH2NL* have been implicated in the expansion of the human cortex (Fiddes et al., 2018; Florio et al., 2016; Heide et al., 2020; Suzuki et al., 2018), supporting predictions that human-specific mutations may influence proliferation of neural progenitor cells during development (Kriegstein et al., 2006; Rakic, 1995). Nonetheless, connecting individual candidate mutations to evolved human traits remains challenging because the large majority of mutations are neutral or low effect size, analyses are low throughput, and we lack a detailed understanding of the divergence in cellular and developmental phenotypes that ultimately give rise to species differences.

In parallel, high-throughput genomics-based approaches have described gene regulatory changes that may contribute to species differences. Because ape primary tissue is largely inaccessible during early development, recent studies have employed stem cell derived models as an experimentally tractable system for comparative analyses of species differences during development. Thousands of cell-type specific gene expression differences have been identified in pluripotent stem cells (Gallego Romero et al., 2015; Marchetto et al., 2013), cardiomyocytes (Pavlovic et al., 2018), endoderm (Blake et al., 2018), neural crest (Prescott et al., 2015), and cortical neurons (Kanton et al., 2019; Marchetto et al., 2019; Pollen et al., 2019). However, these gene expression differences comprise a mixture of neutral changes, causal changes, and indirect downstream consequences and genes that mediate species differences may have conserved expression. Therefore, it can be difficult to ascertain which molecular changes, among hundreds or thousands, drive differences in cellular physiology.

The history of developmental genetics provides a rich template for linking the function of individual genes to organismal phenotypes. Early mutagenesis screens in *Drosophila melanogaster* identified genes critical for body axis patterning (Nüsslein-Volhard and Wieschaus, 1980; Wieschaus and Nüsslein-Volhard, 2016). Many of these genes belonged to highly conserved cell signaling pathways that also coordinate development in vertebrates, such as Wnt (Miller, 2001; Peifer and Wieschaus, 1990), Hedgehog (Hooper and Scott, 1989), and BMP (Costa et al., 1994). More recent efforts in organismal screening involve several international consortia that have generated large collections of knockout mice to investigate more complex vertebrate phenotypes (Dickinson et al., 2016; Hrabě de Angelis et al., 2015; White et al., 2013). The success of these genetic approaches has resulted in the functional annotation of many of the genes and regulatory networks that guide mammalian development. Yet, although many core developmental principles are conserved from fruit flies to mice to humans, these shared molecular functions do not account for how our species evolved to be different.

To apply functional genomics approaches to questions of species divergence, we leveraged recent advances in CRISPR-based technologies that have enabled genome-scale perturbation screens across thousands of human cell lines (Gilbert et al., 2014; Sanjana et al., 2014; Wang et al., 2014). These efforts have mapped landscapes of genetic dependencies with an enrichment of essential genes in coherent pathways that typically cluster by cell type of origin (Pacini et al., 2021; Tsherniak et al., 2017). Extending this approach to studies of comparative evolution might reveal genes or cellular processes with divergent functional roles in homologous cell types. Illuminating the extent and identity of recently-evolved genetic dependencies would complement individual candidate gene approaches, descriptive comparative genomics analyses, and single species loss-of-function studies. However, whether genetic dependencies diverged in closely-related hominin species and how this knowledge could reveal previously unappreciated differences in cellular physiology remains unexplored.

To evaluate the extent of conservation and divergence in genetic dependencies between human and chimpanzee, we established a comparative loss-of-function screening approach in pluripotent stem cells (PSCs). PSCs are a model for the earliest stages of development, capturing features of the inner cell mass of the blastocyst, including the capacity to differentiate into all germ layers at a stage that precedes species differences in developmental timing and cell type composition. The state of pluripotency is well conserved between human and chimpanzee PSCs at the level of the transcriptome, epigenome, and cell fate potential (Gallego Romero et al., 2015), and thus provides a homologous cell type for species comparison. In addition PSCs have greater levels of open chromatin and gene expression than somatic cells (Gaspar-Maia et al., 2011), enabling large-scale study of gene function for genes later expressed in diverse cell types. As PSCs are poised to self-renew or differentiate into all germ layers based on environmental cues, we reasoned that changes in proliferation in PSCs could provide a sensitive measure for species-specific responses to a wide range of genetic perturbations.

Performing genetic screens using an *in vitro* model confers several advantages that could support isolation of molecular and cellular species differences. First, the ability to grow large numbers of PSCs enables a pooled library approach with multiple redundant library elements targeting each gene. Second, laboratory cell culture provides a well-defined and highly controlled environment, which minimizes extrinsic sources of variation. Lastly, the scalability of pooled screening allows for retesting of each cellular phenotype in PSCs derived from multiple individuals of each species to account for individual variation within a species. Thus, we harnessed the power of modern functional genomics to conduct genome-wide CRISPR-interference (CRISPRi) screens in human and chimpanzee PSCs. Despite high levels of conservation, our screens revealed that genetic dependencies can diverge in remarkably short evolutionary time scales, that species differences are organized into coherent pathways and protein complexes, and that human-specific changes have evolved in gene networks promoting G1/S progression in PSCs and neural progenitor cells. In addition to these specific insights, our study establishes a novel and broadly applicable experimental approach for uncovering latent molecular differences between closely-related species.

### Genome-wide CRISPRi screening in human and chimpanzee stem cells

To enable comparative CRISPR-based genetic screening, we engineered CRISPRi machinery (Gilbert et al., 2014) at the CLYBL safe harbor locus (Cerbini et al., 2015) in two human and two chimpanzee pluripotent stem cell lines (Figure 1A). For the two human representatives, we chose two widely used and well-established lines, WTC11 (Kreitzer et al., 2013; Liu et al., 2017; Tian et al., 2019), an induced pluripotent stem cell (iPSC) line, and H1 (Thomson et al., 1998), an embryonic stem cell (ESC) line. For the chimpanzee representatives, we chose two robust iPSC lines used in previous studies: C3649 and Pt5-C (Gallego Romero et al., 2015; Ryu et al., 2018) (Table S1).

**Figure 1.**
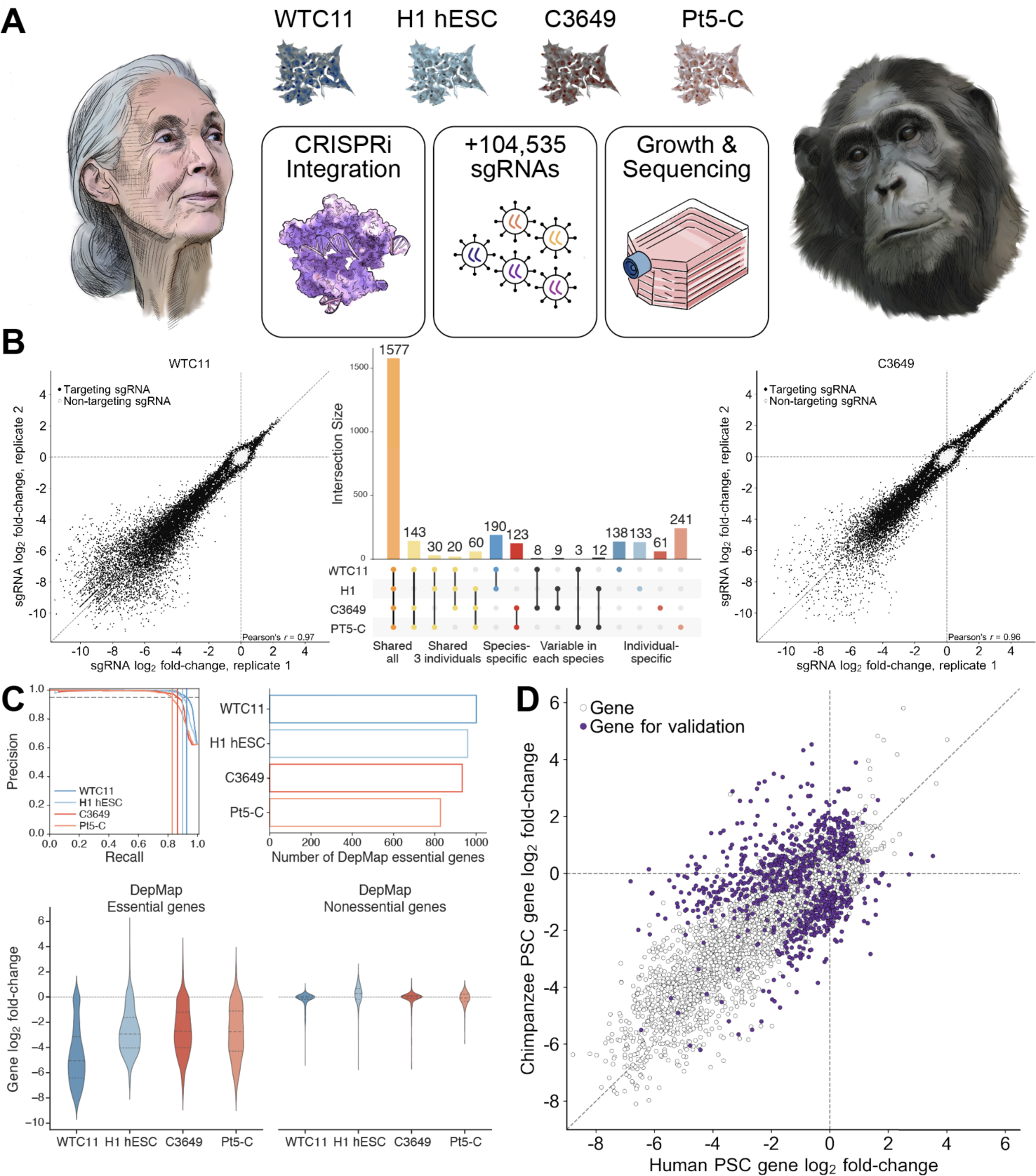
Genome-wide CRISPRi screens in human and chimpanzee stem cells identify candidate species-specific genetic dependencies. (A) Schematic of CRISPRi screening approach. Two human (WTC11 and H1) and two chimpanzee (C3649 and Pt5-C) PSC lines were engineered to express dCas9-KRAB, infected with the lentiviral hCRISPRi-2 sgRNA library, and grown competitively for 10 days. Depleted and enriched sgRNAs were detected by high-throughput sequencing. (B) Scatterplots of sgRNA fold-change for WTC11 and C3649 technical replicates and UpSet plot showing the intersection of essential genes across screens. (C) Precision-recall analysis (top left) for each screen. Precision and recall were determined using DepMap essential and nonessential genes. The number of DepMap essential genes (top right) identified by MAGeCK (5% FDR, log2 fold-change < −1.5). Distribution of fold-change for DepMap essential (bottom left) and nonessential (bottom right) genes. (D) Species-level gene fold-change across genome-wide CRISPRi screens. Gene-level phenotypes were computed as the mean of the three sgRNAs with the largest absolute fold-change. sgRNAs lacking perfect-match targets in the chimpanzee genome were excluded from analysis.

To identify genes that modify cellular growth and survival, we infected each cell line with the genome-wide lentiviral hCRISPRi-v2 sgRNA library (Horlbeck et al., 2016, Table S2) (5 sgRNAs/gene), selected for sgRNA-expressing cells with puromycin, cultured cells for 10 days, and quantified sgRNA enrichment and depletion by high-throughput sequencing (Table S3). While hCRISPRi-v2 was designed to target the human genome, 77.4% of sgRNAs perfectly matched targets in the chimpanzee reference genome (panTro6); sgRNAs with mismatches were not considered for analyses of species differences (Figure S1A, STAR methods). Across all four screens, we observed robust depletion of sgRNAs targeting common essential genes and enrichment of sgRNAs targeting proliferation-suppressor genes. Analysis of technical and biological replicates revealed strong sgRNA correlations for replicates of the same cell line (Pearson’s *r* = 0.80 to 0.97, Figure 1B) and for different cell lines within species (*r* = 0.69 to 0.83). In addition, all four genetic screens sensitively and precisely distinguished Dependency Map (DepMap) common essential and nonessential genes (Blomen et al., 2015; Hart et al., 2014), recalling 82.9% to 92.6% of common essential genes at 95% precision (Figure 1C).

We next sought to identify genes with species-specific effects on cellular proliferation. To do so, we utilized MAGeCK (Li et al., 2014) and developed a bootstrapping-based method that accounted for both the number of significantly enriched or depleted sgRNAs targeting a gene and the magnitude of sgRNA log2 fold-change (Figures S1B-D, STAR methods). While the large majority of essential genes were shared between species (Figure 1B), we identified 583 candidate species-specific essential genes and 202 candidate species-specific proliferation-suppressor genes (Figure 1D). Importantly, this approach identified far fewer candidate genes exclusively shared between one individual of each species (*n* = 3 to 12 genes, Figure 1B), highlighting the influence of species on gene essentiality. These results establish a CRISPRi-based approach that overcomes both technical noise in genome-scale screening and variability between PSC lines to reveal candidate species differences.

### Human and chimpanzee can be distinguished by genetic dependencies

We next sought to validate candidate species differences across multiple independently-derived human and chimpanzee PSCs to distinguish species differences from those driven by individual variation (Cahan and Daley, 2013; Kilpinen et al., 2017; Prado-Martinez et al., 2013), adaptation to cell culture (Baker et al., 2007), or somatic cell reprogramming (Merkle et al., 2017). We engineered new CRISPRi stem cell lines from four human (H20961B, H21792A, H23555A, H28126B) and four chimpanzee (C3624K, C8861G, C40280L, C40290F) individuals. To minimize technical variation, we selected cell lines with normal karyotypes that were reprogrammed with identical protocols (Table S1) and maintained in identical media. Furthermore, cell lines from both species were previously shown to differentiate into all three germ layers via teratoma formation and embryoid body assays, functionally validating pluripotency (Gallego Romero et al., 2015). Finally, the human and chimpanzee lines shared comparable pluripotency scores with strongly overlapping patterns of H3K27me3 and H3K27ac at pluripotency genes (Gallego Romero et al., 2015) and similar transcriptional trajectories of differentiation (Blake et al., 2018), suggesting the lines are in a comparable state of pluripotency. We analyzed copy number variation using CaSpER (Serin Harmanci et al., 2020) and genome sequencing coverage to rule out the presence of large duplications or deletions (Figure S2). In addition, we assessed CRISPRi cell lines for p53-responsiveness by measuring sensitivity to nutlin-3a, a small molecule MDM2 inhibitor that induces p53-dependent autophagy and apoptosis (Setoguchi et al., 2016; Vassilev et al., 2004). All 11 new human and chimpanzee lines plus WTC11 and H1 were MDM2/p53-responsive, while C3649 and Pt5-C, the chimpanzee lines used for genome-scale screening, were nonresponsive to MDM2/p53 perturbations (Figures S1E and S1F).

To enable secondary screening, we designed a comparative essential validation (CEV-v1) library consisting of 7,847 sgRNAs targeting the transcriptional start sites of 963 genes from our genome-scale datasets (8 sgRNAs/gene, STAR methods) and 1,845 negative-control sgRNAs (Figure 2A; Figure S3, Table S4). Due to the scalability of pooled screening, we targeted an inclusive set of genes with significant or suggestive differences between species in the primary screens as well as gene families with notable evolutionary histories (Dennis and Eichler, 2016; Pontis et al., 2019). To reduce human-specific bias, we required that every sgRNA in CEV-v1 perfectly match target sites in the human (hg38) and chimpanzee (panTro6) reference genomes (McKenna and Shendure, 2018). In total, we performed 16 CRISPRi screens the CEV-v1 sgRNA library (Figure 2A, Table S5). The validation screens were performed in the four newly constructed CRIPSRi PSC lines of each species. In addition, we retested three of the four PSC lines used for genome-scale screening (Pearson’s *r* = 0.76 to 0.89) and performed biological replicate screens in separate laboratories for five cell lines (3 human lines, 2 chimpanzee lines, *r* = 0.70 to 0.92). Notably, hierarchical clustering of cell lines by the similarity of their sgRNA profiles separated all the human (including ESCs and iPSCs) from all the chimpanzee individuals (Figure 2B). Decomposition of each cell line’s sgRNA profile by principal component analysis also grouped individuals by species, with the main axes of variation relating to shared changes in sgRNA representation over time (PC1) and species-specific changes (PC2) (Figure 2C). Together, our findings show that stem cells from humans and chimpanzees can be distinguished by their responses to genetic perturbations.

**Figure 2.**
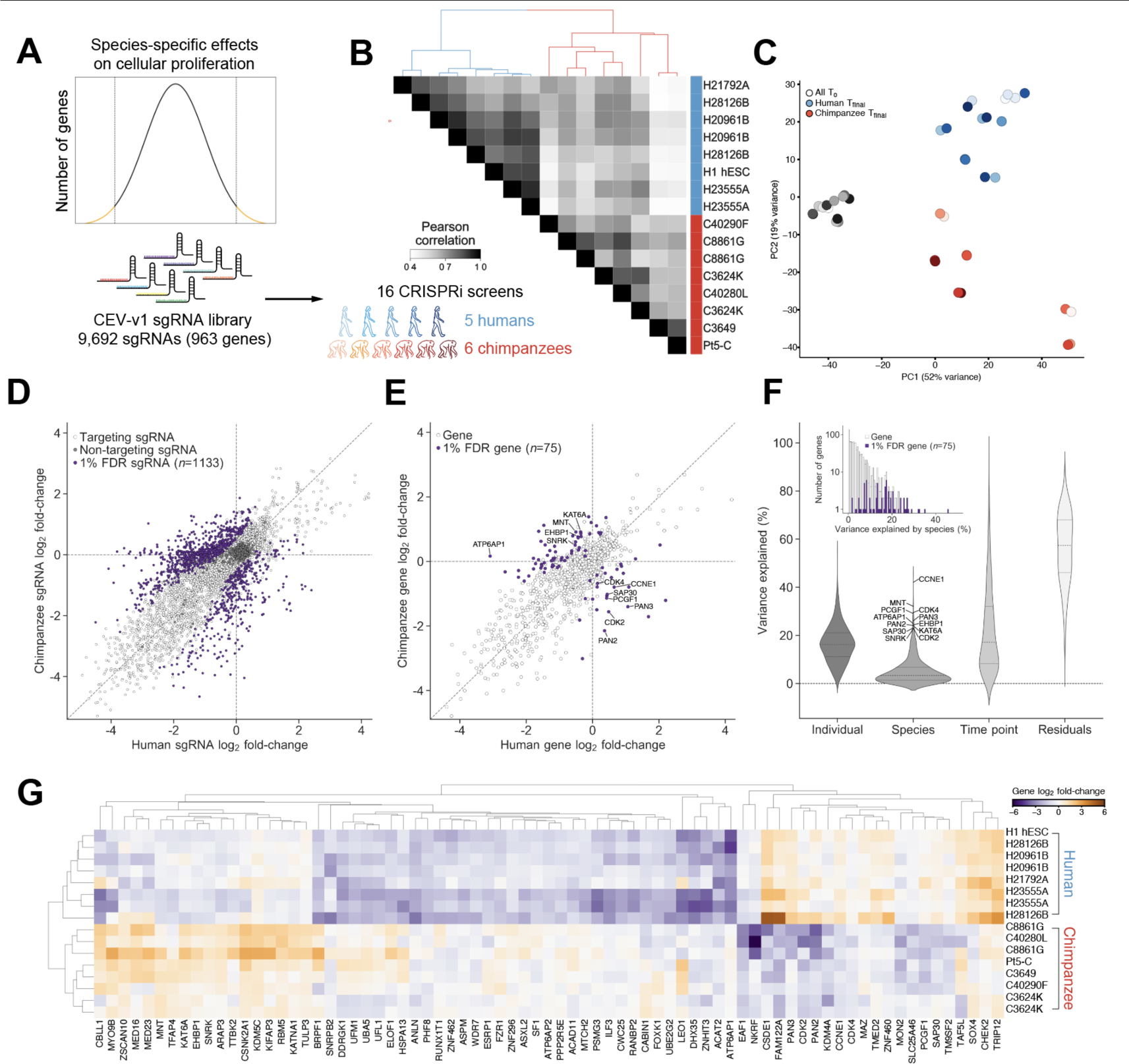
Species-specific genetic dependencies validate across five human and six chimpanzee individuals. (A) Schematic of validation sgRNA library design and CRISPRi screening approach. (B) Heatmap of Pearson correlations and hierarchical clustering for sgRNA profiles across 16 validation CRISPRi screens. Individuals listed twice are replicate screens performed in separate laboratories. (C) Principal component analysis of sgRNA counts at t0 (black circle) and tfinal (red and blue circles). (D) Scatterplot of log2 fold-change of sgRNA counts, weighted and normalized by DESeq2. 1,133 sgRNAs with significant species differences (FDR < 0.01) colored in purple and negative-control sgRNAs colored in dark gray. (E) Species-level gene fold-change across validation CRISPRi screens. Gene-level phenotypes were computed as the mean of the four sgRNAs with the largest absolute fold-change. The 12 genes with the greatest variance in sgRNA fold-change attributable to species are labeled. (F) Dream-variancePartition analysis for quantifying sources of variation in sgRNA counts attributable to individual, species, and timepoint (t0 vs. tfinal). (G) Heatmap of gene fold-change and hierarchical clustering for 75 genes with species-specific effects on cellular proliferation across validation CRISPRi screens (1% FDR).

### Molecular nature of core species-specific genetic dependencies

We next sought to identify genes underlying the differences between human and chimpanzee sgRNA profiles. Using generalized linear models borrowed from DE-seq2 (Love et al., 2014) we identified 1,133 sgRNAs with evidence for differences between species (1% false discovery rate (FDR), |chimpanzee−human log2 fold-change| ≥ 0.5), while negative-control sgRNAs were tightly distributed around zero (Figure 2D). Using α-RRA (Li et al., 2014) to combine sgRNA *P*-values, we found 75 genes with robust species-specific effects on cellular proliferation at a 1% FDR (Figure 2E-G; STAR methods). This substantial reduction in the number of significant genes from the CEV-v1 candidate gene pool was the result of a combination of factors: 1) the use of more stringent statistical thresholds, 2) a fraction of genes that replicated in validation screens of the original cell lines but not in the additional cell lines 3) the exclusion of genes whose effects on cellular proliferation depended on *TP53* status. Together, these findings reveal a stringent set of species-specific genetic dependencies that emerged in recent human and chimpanzee evolution.

We first explored whether species-specific genetic dependencies could relate to changes in the coding sequence or regulation of the target genes themselves that might suggest new species-specific activities of these genes. Several genes in the set exhibited unexpected coding sequence changes. For example, ASPM, which causes microcephaly when mutated, contains protein domains with signatures of positive selection in the human lineage (Evans et al., 2004; Kouprina et al., 2004; Zhang, 2003) and was essential in human but not chimpanzee PSCs. Similarly, KATNA1, which physically interacts with ASPM to promote microtubule disassembly at mitotic spindle poles (Jiang et al., 2017), contains a nearly fixed modern human-specific mutation that is distinct from the Neanderthal and chimpanzee allele (Kuhlwilm and Boeckx, 2019) and acted as a suppressor of proliferation in chimpanzee but not human cells. However, these examples were exceptions, and signatures of adaptive selection, as well as overall non-synonymous substitutions, were depleted among the set of 75 species-specific genetic dependencies compared to the genome wide distribution (p < 0.01, p < 10^-6, respectively, Kolmogorov–Smirnov test, Figure S4A-B). Several genes with divergent genetic dependencies also displayed quantitative gene expression changes. For example, *MTCH2*, a gene involved in mitochondrial metabolism and apoptosis (Gross, 2016) displayed significantly higher expression in human PSCs (fold change = 1.28, FDR < 10^-3^) and was specifically essential in human PSCs. In contrast, *ACAT2*, a gene involved in lipid metabolism, exhibited significantly higher expression in chimpanzee PSCs (fold change = 2.73, FDR < 10^-6^), but was also specifically essential in human cells. Despite these examples, the 75 gene set was also depleted for species differences in gene expression (*P* = 0.035, Figure S4C). Together, these analyses suggest that coding or regulatory changes do not account for the majority of species differences and that a multitude of indirect effects may underlie divergent dependencies (Liu et al., 2019b).

We next asked whether species-specific genetic dependencies involved groups of genes known to interact, a pattern that could suggest divergent requirements for conserved pathways. As essentiality phenotypes are typically shared among genes within known functional modules (Kim et al., 2022; Wainberg et al., 2021), such coherence could also provide an additional test of internal consistency. Indeed, functionally related genes emerged with consistent patterns of depletion or enrichment within each species. Analysis using the STRING database (Szklarczyk et al., 2021) revealed an enrichment for protein-protein interactions (p < 10^-6^) and components of several well established biological processes (Figure 3A). For example, we observed that all five core components of the UFMylation pathway (*UFM1*, *UFL1, UFC1*, *UBA5*, *DDRGK1*) were essential only in human PSCs (Figure S4D). By contrast, all four subunits of the MOZ histone acetyltransferase complex (*KAT6A*, *BRPF1*, *ING5*, *EAF6*) acted as proliferation suppressors in chimpanzee PSCs (Figure S4D). Accessory proteins to the vacuolar-type ATPase (*ATP6AP1*, *ATP6AP2*) and the highest ranking DepMap co-dependent gene *WDR7* (Tsherniak et al., 2017), were specifically essential in human PSCs, whereas core subunits were essential in both species (Figure 3B). Strikingly, human PSCs were robust to depletion of cell cycle regulators cyclin-dependent kinase 2 (*CDK2*), its activating partner *Cyclin E1* (*CCNE1*), and cyclin-dependent kinase 4 (*CDK4*). For all three genes, we observed at least six sgRNAs that were essential across six chimpanzee individuals but nonessential across six human individuals (Figure 3C).

**Figure 3.**
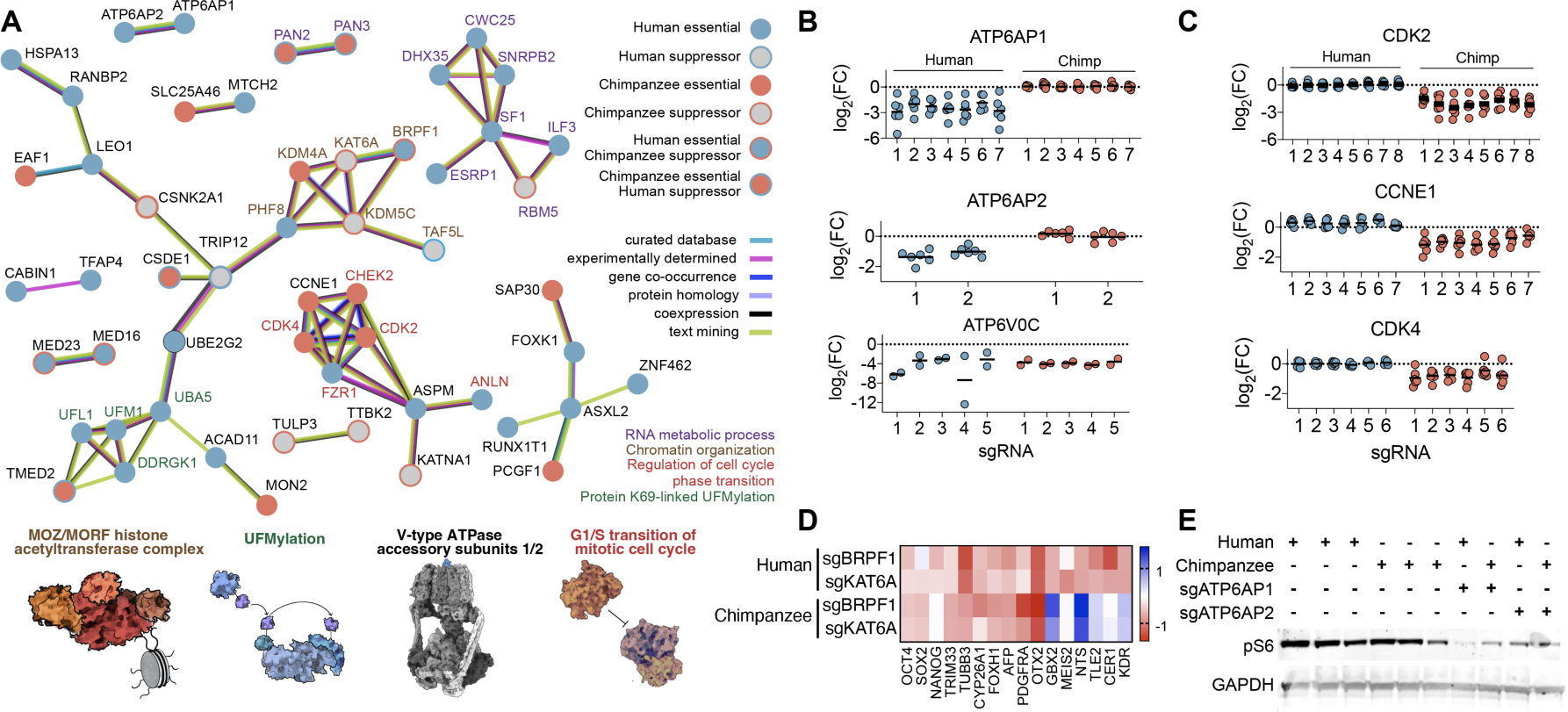
Core species-specific genetic dependencies. (A) Species-specific genetic dependencies with STRING protein-protein associations. Illustrations of pathways and protein complexes with coherent species-specific effects. (B) Strip plots of fold-change for sgRNAs targeting *ATP6AP1*, *ATP6AP2* and *ATP6V0C* (C) *CDK2*, *CCNE1*, and *CDK4*. Each circle represents the sgRNA fold-change for one sgRNA in one human (blue) or chimpanzee (red) individual. Each stripplot contains a variable number of columns, corresponding to the number of significant sgRNAs targeting each gene. (D) Heatmap of RNA-seq expression data for human (WTC11) and chimpanzee (C3649) cells depleted for KAT6A or BRPF1. (E) Western blot for phospho-S6 expression and GAPDH loading control for three wild-type human (H1, 21792A, and 28128B) and three wild-type chimpanzee (3624K, 40280L, and 8861G) cell lines, and cell lines depleted for ATP6AP1 or ATP6AP2 (28128B and 40280L).

The consistent depletion of many sgRNAs targeting the same gene and multiple genes involved in the same biological process indicates that off-target activity is unlikely to explain proliferation differences between species. In principle, species-specific differences in our CRISPRi screens could also result from differential effectiveness of CRISPRi-mediated transcriptional repression (e.g., due to histone occupancy or transcriptional start site variability). To evaluate this possibility, we measured the efficacy of sgRNA-mediated repression for several candidate genes (*CDK2*, *CCNE1*, and *RBL1*). In all cases, measurements of transcript abundance by qRT-PCR revealed >90% knockdown in both species (Figure 4D, Table S6), suggesting that proliferation differences were not driven by incomplete knockdown efficiency in one species. In summary, these results highlight the ability of our screening approach to isolate biologically meaningful networks of genes that mediate species differences in cell behavior when perturbed, in contrast to gene expression profiling, which often reveals a complex mixture of both direct and indirect effects.

**Figure 4.**
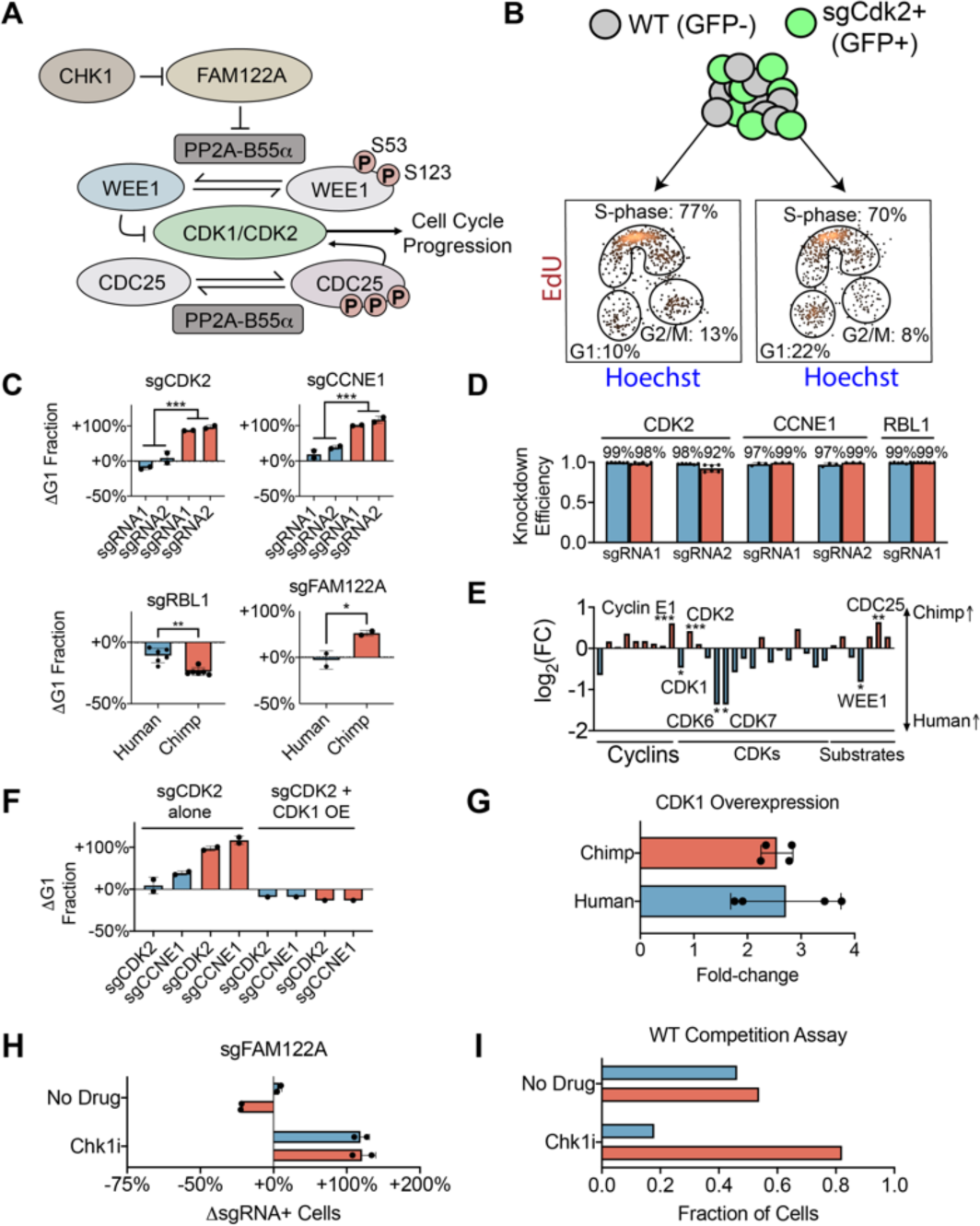
Divergent regulation of cell cycle progression in human and chimpanzee cells. (A) Schematic for Cdk1/Cdk2 regulatory network. Cdk1 and Cdk2 phosphorylate key substrates Wee1 and Cdc25, leading to degradation of Wee1 and activation of Cdc25. Phosphatase PP2A dephosphorylates Wee1 and Cdc25 at the same sites. Chk1 inhibits FAM122A, and FAM122A inhibits PP2A. (B) Cell cycle proportions in chimpanzee wild-type cells (GFP-) and sgRNA containing cells (GFP+) grown in co-culture. (C) Change in the fraction of cells in G1 phase upon knockdown of *CDK2* (*P* < 10^-3^), *Cyclin E1* (*P* < 10^-3^), *RBL1* (*P* < 10^-2^), and *FAM122A* (*P* < 0.05). (D) qRT-PCR measurements of knockdown efficiency in human (blue) and chimpanzee (red) PSCs. (E) Comparative gene expression data from human and chimpanzee PSCs for core cell cycle regulators (* *P* < 0.05, ** *P* < 10^-2^, *** *P* < 10^-3^). (F) Change in the fraction of cells in G1 phase upon overexpression of *CDK1* in conjunction with *CDK2* or *Cyclin E1* knockdown. (G) qRT-PCR measurements of the degree of *CDK1* overexpression. (H) Change in the fraction of FAM122A sgRNA containing cells in the presence of no drug or Chk1 inhibitor prexasertib (Chk1i). (I) Fraction of wild-type human (blue) vs. wild-type chimpanzee (red) cells grown in co-culture in the presence of no drug or Chk1i.

### Genetic perturbations elicit divergent cellular responses

While both our primary screen and validation screens measured growth and survival, changes in proliferation can reflect a wide range of cellular phenotypes, from differentiation to growth factor signaling. We first investigated the growth advantage of chimpanzee cells depleted for components of the MOZ histone acetyltransferase complex. MOZ acetylates histone H3 at lysines 9 and 14, and knockout mice are embryonic lethal at E14.5 due to defects in hematopoiesis (Katsumoto et al., 2006; Ullah et al., 2008; Yang, 2015). Given the role of the MOZ complex in epigenetic modifications, we hypothesized that there would be species differences in gene expression response. We performed transcriptional profiling of human and chimpanzee cells infected with sgRNAs targeting *BRPF1* or *KAT6A*, normalized to wild-type cells of the same PSC line (Figure 3D, Table S7). Expression changes were strongly correlated between the two perturbations with Pearson’s correlation coefficient 0.80 in human cells and 0.83 in chimpanzee cells. Gene ontology enrichment analyses of expression changes combined across both sgRNA perturbations revealed that human cells upregulated genes involved in TGF-beta signaling, cell differentiation, and morphogen activity upon MOZ depletion (Figure S5). In contrast, chimpanzee cells overexpressed genes involved in mTORC1 signaling and Myc targets, consistent with their species-specific growth advantage. In both species, we observed a transcriptional signature consistent with heterogeneous loss of pluripotency. Bulk expression levels of key pluripotency genes *OCT4*, *SOX2*, and *NANOG* were largely unchanged or slightly higher in sgRNA containing cells. However, a broad range of marker genes for multiple germ layers were upregulated. Both species upregulated markers associated with primary ectoderm (*TRIM33*, *TUBB3*), endoderm (*AFP*, *FOXH1*, *CYP26A1*), and mixed lineages (*PDGFRA*, *OTX2*) (Maguire et al., 2013). However, human cells selectively upregulated additional markers of ectoderm (*GBX2*, *MEIS2*), endoderm (*NTS*, *TLE2*, *CER1*), and mixed lineages (*KDR*) compared to chimpanzee cells. Thus, we propose that the MOZ complex plays a conserved role in maintaining an epigenetic barrier to differentiation for both species (Ciceri et al., 2022). However, upon loss of MOZ-mediated histone acetylation, human cells activate expression of a wider range of differentiation markers, suggesting that species differences in epigenetic potential may exist pluripotent cells.

We next investigated the human-specific sensitivity to loss of *ATP6AP1* and *ATP6AP2*. ATP6AP1 and ATP6AP2 are accessory proteins to the lysosomal V-type ATPase. As the main proton pump responsible for maintaining the pH gradient of the lysosome, non-duplicated core subunits of the V-ATPase were essential in both species, as expected, (Figure 3B). Cryo-electron microscopy of the V-ATPase complex has implicated ATP6AP1 in the assembly of the V0 complex of the V-ATPase (Wang et al., 2020). In addition, ATP6AP1 is comprised of a transmembrane helix and an extensive luminal domain that bears extensive structural homology to lysosomal-associated membrane proteins (LAMPs) and forms extensive contacts with ATP6AP2. Staining with LysoTracker Red and LysoSensor green in *ATP6AP1* depleted cells revealed no significant defects in maintenance of lysosomal pH (Figure S6). These results are consistent with the core function of V-ATPase being strictly essential. However, loss of *ATP6AP1* has also been implicated in major cellular signaling pathways that are mediated by the lysosome (Jewell et al., 2013; Zhou et al., 2009; Zoncu et al., 2011). Thus, we performed a western blot to measure phosphorylation of ribosomal protein RPS6 (pS6), a well-established downstream effector of mTORC1 activity. Depletion of *ATP6AP1* or *ATP6AP2* resulted in diminished pS6 signal in both species. However, pS6 was selectively abolished in human cells depleted for *ATP6AP1* (Figure 3E). These data thus link the human-specific growth defect of *ATP6AP1* sgRNAs observed in our pooled screens to an increased reliance of human cells on ATP6AP1-mediated mTORC1 signaling.

### Human PSCs are robust to depletion of CDK2 and Cyclin E

Based on the enrichment of genes that regulate cell cycle progression among species-specific genetic dependencies, we next connected the growth phenotypes observed upon depletion of cell cycle factors to more precise cell cycle defects (Figure 4A). To do so, we measured the proportion of cells in G1, S-phase and G2/M via incorporation of the thymidine analogue 5-ethynyl-2′-deoxyuridine (EdU) and Hoechst, a DNA-binding dye. Consistent with the early mammalian embryo, PSCs undergo rapid cell cycle progression with a shortened G1 phase compared to somatic cells (Becker et al., 2006). For wild-type cells, only ∼10% of cells were classified in G1 phase (Figure 4B). However, the absolute fraction of G1 cells was influenced by environmental factors such as confluence and nutrient availability. Therefore, we measured the effect of CRISPRi-mediated gene repression in an internally controlled co-culture, with wild-type cells (GFP-) and sgRNA-expressing cells (GFP+) mixed within the same well. Knockdown of *CDK2* or *Cyclin E1* in chimpanzee PSCs led to a roughly two-fold accumulation of cells in G1 (Figure 4C; *P* < 10^-3^ for both), consistent with the well-established role of Cyclin E1-Cdk2 in regulating the G1/S transition. By contrast, knockdown of *CDK2* in human PSCs had no effect on cell cycle progression, and knockdown of *Cyclin E1* produced only a limited accumulation of G1 cells. We confirmed that these differences were not mediated by incomplete sgRNA-mediated knockdown in human PSCs (Figure 4D). Thus, our data suggest that human PSCs are less dependent on Cyclin E1-Cdk2 for G1/S phase transition than chimpanzee PSCs.

Cyclin E1-Cdk2 is a central regulator of the G1/S cell cycle transition (Hochegger et al., 2008) and is commonly essential (Pacini et al., 2021) and frequently dysregulated across human cancer cell lines (Hwang and Clurman, 2005). In contrast, *CDK2*^-/-^ knockout mice are fully viable and develop normally, though with reduced body size (Berthet et al., 2003; Ortega et al., 2003). Subsequent studies showed that cell cycle progression could be rescued in the absence of Cyclin E1-Cdk2 by Cyclin A-Cdk1 and Cyclin E-Cdk1 activity (Aleem et al., 2005; Satyanarayana and Kaldis, 2009). Therefore, we reasoned that human cells might compensate for the loss of Cyclin E1-Cdk2 via stronger Cyclin A2-Cdk1 activity, as cyclin homologs Cyclin E2 and Cyclin A1 are not expressed in PSCs. Consistent with this model, *CDK1* was more highly expressed in human PSCs (FDR < 10^-2^), while *CDK2* and *Cyclin E1* were more highly expressed in chimpanzee PSCs (FDR < 10^-3^) (Figure 4E; Figure S7). As a functional test, we overexpressed *CDK1* in chimpanzee PSCs in conjunction with sgRNA-mediated repression of *CDK2* or *Cyclin E1* and quantified the progression of cells thru G1 phase. We found that 2.5-fold overexpression of *CDK1* was sufficient for rescuing the sensitivity of chimpanzee PSCs to *CDK2* or *Cyclin E1* depletion and accelerated progression through G1/S phase transition (Figure 4F-G).

Next, we extended our co-culture studies to additional cell cycle regulators with known interactions with Cyclin E-Cdk2. Given the dependence of chimpanzee PSCs on cyclin E-Cdk2, we reasoned that repression of an inhibitor of this cyclin-Cdk complex might have species-specific effects on cell cycle progression. We first investigated the consequences of repressing retinoblastoma-like 1 (*RBL1*/p107), a tumor suppressor homologous to retinoblastoma protein (*RB*). Rbl1, like Rb, classically represses cell cycle via inhibition of E2F transcription factors (Rubin et al., 2020; Shirodkar et al., 1992; Zatulovskiy et al., 2020). However, E2F is de-repressed in rapidly-dividing stem cells compared to other cell types due to the need for rapid cell cycling (Liu et al., 2019a). Rbl1 also possesses an ability unique among Rb family proteins to directly inhibit the kinase activities of cyclin A/E-Cdk2 (Zhu et al., 1995). Accordingly, we observed that repression of *RBL1* but not *RB* resulted in faster growth and a reduction in the fraction of cells in G1 in both species (Figure 4C). However, *RBL1* effects were larger for chimpanzee cells (P < 0.01). Given the accumulation of chimpanzee PSCs in G1 upon repression of *Cyclin E* or *CDK2* (Figure 4C), these results further support a model in which cyclin E-Cdk2 exerts greater control over G1/S transition in chimpanzee compared to human PSCs.

To further link regulation of Cdk1/2 to species divergence in cell cycle progression, we focused on family with sequence similarity 122A (*FAM122A*) (Liu et al., 2021; Zhou et al., 2020), a chimpanzee-specific essential gene. FAM122A acts as an inhibitor phosphatase PP2A-B55α (Fan et al., 2016), which in turn acts in opposition to Cdk1 and Cdk2 by dephosphorylating key substrates such as Wee1 and Cdc25 (Mochida et al., 2009) (Figure 3A). We observed that loss of *FAM122A* phenocopied loss of *CDK2* and led to accumulation of G1 cells in chimpanzee PSCs, but not in human (Figure 4C; *P* < 0.05). Our results suggest that higher endogenous *CDK1* levels could overcome *FAM122A* loss and PP2A phosphatase activation to promote cell cycle re-entry in human PSCs. However, the balance of kinase and phosphatase activity was more sensitive to perturbation in chimpanzee cells, as *CDK2* repression or activation of PP2A led to a delayed G1/S transition.

Finally, we further examined species differences in PSC cell cycle progression using pharmacological approaches. Based on prior literature (Li et al., 2020), we tested the interaction between *FAM122A* and prexasertib, a Chk1 inhibitor (CHK1i) currently undergoing phase II oncology trials (Byers et al., 2021; Gatti-Mays et al., 2020; Lampert et al., 2020). As with cancer cells, the rapid cell divisions in PSCs render them sensitive to replication stress and DNA damage, conferring vulnerability to prexasertib. As expected, *FAM122A* knockdown PSCs of both species were resistant to prexasertib (Figures 4H; Li et al., 2020). However, with no sgRNA present, wild-type human PSCs were more acutely sensitive to Chk1 inhibition compared to chimpanzee (Figure 4I). These data suggest that chimpanzee PSCs potentially enforce a more robust S phase and G2/M checkpoint compared to human PSCs.

### Cell cycle perturbations alter neural progenitor cell expansion

We wondered whether the molecular differences that we observed between species would also manifest in differentiated cell types. As differences in G1/S regulation have long been hypothesized as an evolutionary mechanism for changing brain size (Dehay and Kennedy, 2007; Lukaszewicz et al., 2005; Pilaz et al., 2009; Taverna et al., 2014), we investigated whether human-specific robustness to depletion of cell cycle factors would persist in neural progenitor cells (NPCs). Previous studies have established both the necessity and sufficiency of genes promoting G1/S transition for proliferative NPC divisions in animal model systems (Calegari et al., 2005; Lange et al., 2009, 2009; Lim and Kaldis, 2012; Lukaszewicz et al., 2005; Nonaka-Kinoshita et al., 2013; Pilaz et al., 2009). However, it is not known whether human NPCs possess recently evolved characteristics that imbue them with an enhanced ability to maintain proliferative divisions. We generated CRISPRi human and chimpanzee NPCs and assessed how depletion of *Cyclin E1*, *CDK2*, *RBL1*, and *FAM122A* affected cell cycle progression and self-renewal (Figure 5A-D; Figure S8A-C). In contrast to PSCs, NPCs undergo substantially slower progression through cell cycle, with ∼50% of cells in G1 phase compared to ∼10% in PSC (Figure 5B). Nonetheless, knockdown of *Cyclin E1* (*P* < 0.05) or *CDK2* (*P* < 10^-3^) caused an additional accumulation of chimpanzee, but not human, NPCs in G1 (Figure 5C). Meanwhile, *RBL1* knockdown increased NPC proliferation and reduced the fraction of G1 cells in both human and chimpanzee (Figure 5C). Lastly, knockdown of *FAM122A* resulted in G2/M accumulation in chimpanzee but not human NPCs (Figure 5D), implying a greater role for PP2A activation at G2 in chimpanzee NPCs compared to PSCs. Our results suggest that the increased robustness of human NPCs to depletion of regulators of G1/S progression could potentially bias human cells towards prolonged proliferative divisions, as has been proposed by developmental models.

**Figure 5.**
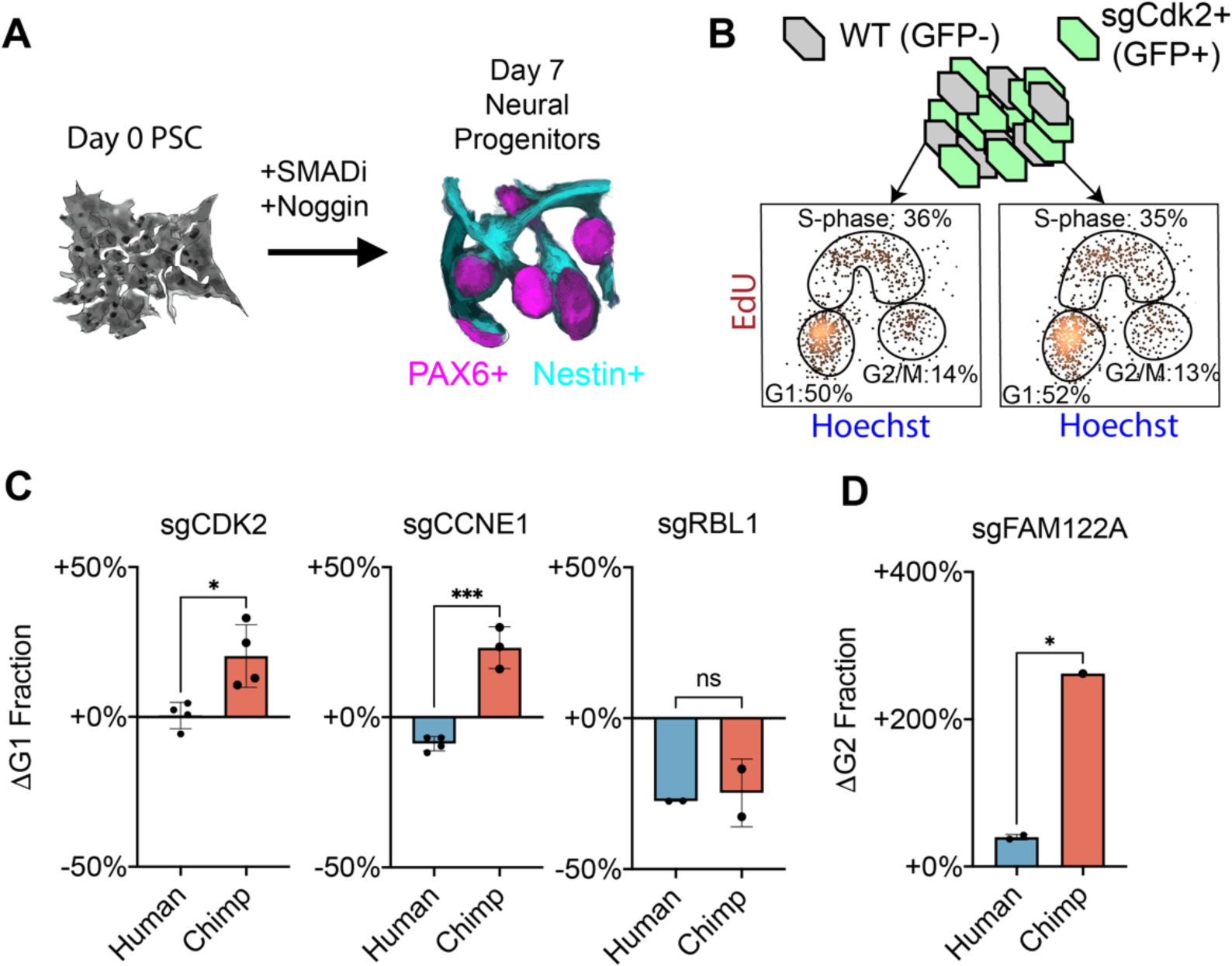
Differentiation into neural progenitor cells. (A) Schematic for differentiation of PSCs into neural progenitor cells (NPCs). (B) Cell cycle proportions in human wild-type neural progenitor cells (GFP-) and sgRNA containing cells (GFP+) grown in co-culture. (C) Change in the fraction of NPCs in G1 phase upon depletion of *CDK2* (*P* < 0.05), *Cyclin E1* (*P* < 10^-3^), or *RBL1* (n.s.). (D) Change in the fraction of NPCs in G2 phase upon depletion of *FAM122A* (*P* < 0.05).

### Evolutionary origin of molecular species-differences

To determine the evolutionary origin of human- and chimpanzee-specific genetic dependencies, we extended our comparative studies to orangutan PSCs (Field et al., 2019). While humans and chimpanzees diverged roughly seven million years ago (Langergraber et al., 2012), orangutans diverged from other great apes 13-18 million years ago (Glazko and Nei, 2003). Thus, we could infer by maximum parsimony that any genetic dependencies shared between orangutans and chimpanzees but not humans were likely to have been present in the common ancestor and subsequently diverged in the human lineage. We performed three-way species comparisons across genes representing several biological processes with coherent species differences in our dataset using sgRNAs with perfectly-matched targets in all three species. For two sgRNAs targeting *CDK2*, we observed a significant depletion of sgRNA-expressing cells over the course of ten days in both chimpanzee and orangutan PSCs (Figure 6A). In contrast, no such depletion was observed in human PSCs. We further confirmed that the differences we observed were not due to differences in sgRNA activity, as knockdown efficiency exceeded 90% in all three species. In addition, we further observed human-specific robustness to repression of *CDK4* (Figure 6B) and *Cyclin E1* (Figures S9A and S9B). Based on these data, we inferred that robustness to perturbations of the G1/S transition evolved along the human lineage, otherwise dependence on *CDK2*, *CDK4*, and *Cyclin E1* would have had to evolve on two separate occasions in the chimpanzee and orangutan lineages with the same direction of effect for each gene. Next, we evaluated the human-specific sensitivity to repression of *ATP6AP1*. We observed that the *ATP6AP1* sensitivity was not shared by chimpanzee or orangutan PSCs, suggesting that altered responses to cellular metabolism, including the increased reliance on ATP6AP1 for mTORC1 signaling that we observed also evolved along the human lineage (Figure 6C).

**Figure 6.**
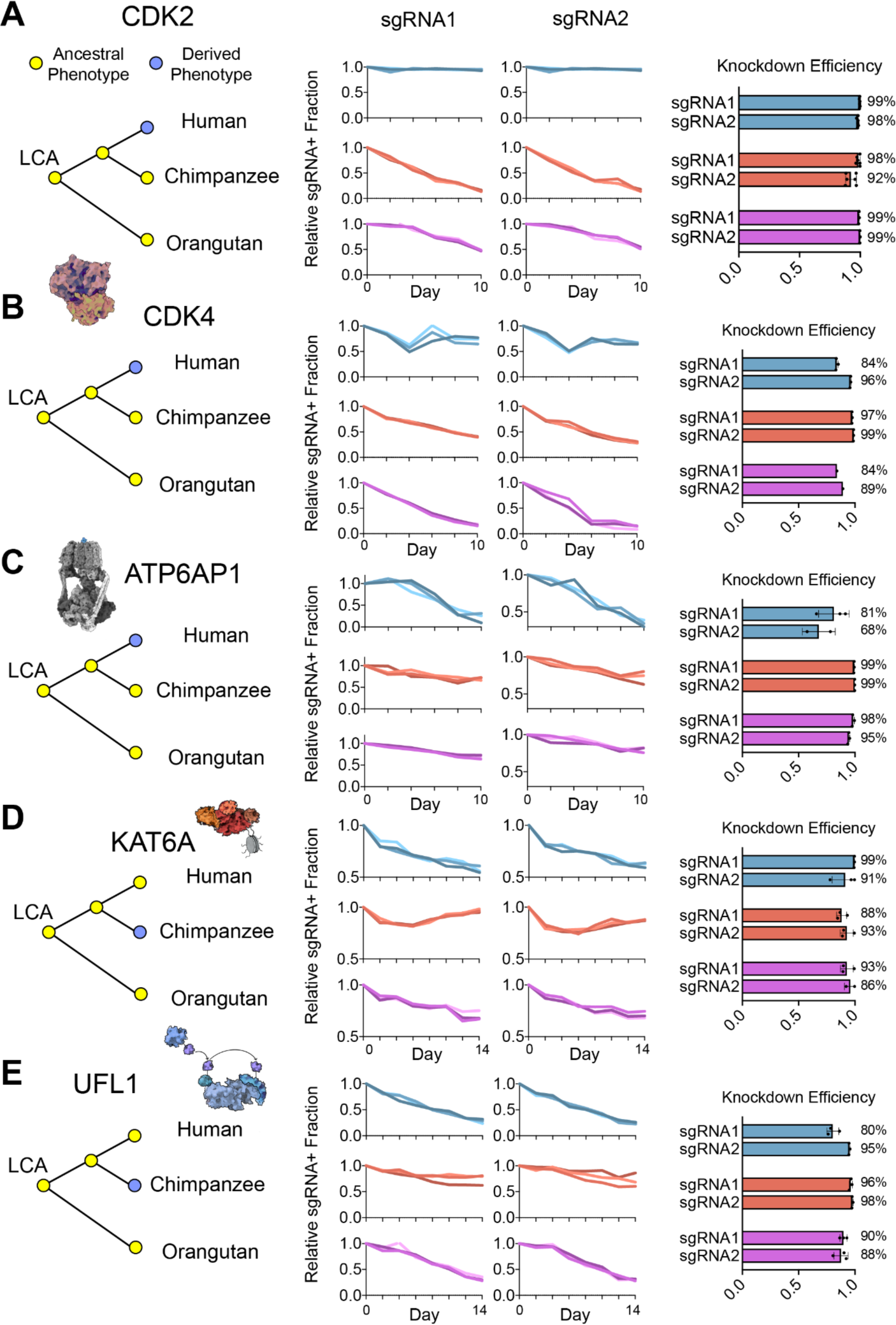
Orangutan PSCs reveal evolutionary origin of species-specific genetic dependencies. (A) Change in the relative fraction of CDK2 sgRNA containing cells over time in human, chimpanzee, and orangutan PSCs. qRT-PCR measurements of sgRNA knockdown efficiency for each sgRNA in all three species. (B-E) Relative sgRNA fraction over time and qRT-PCR measurements for sgRNAs targeting (B) *CDK4*, (C) *ATP6AP1*, (D) *KAT6A*, and (E) *UFL1*.

By contrast, repression of *KAT6A* promoted proliferation in chimpanzee PSCs but not in human or orangutan PSCs, arguing that this molecular feature was derived in chimpanzees (Figure 6D). Similarly, sensitivity to *UFL1* repression was common to human and orangutan PSCs but diverged in chimpanzee PSCs (Figure 6E). The bidirectional polarization of orangutan PSCs towards both human and chimpanzee PSCs, depending on biological pathway, provides an additional line of evidence that the species-specific genetic dependencies that we observed did not arise from two distinct pluripotency states in human and chimpanzee PSCs. If this were the case, the orangutan PSCs should consistently polarize more closely towards one species and not the other. In sum, our data indicate that distinct genetic dependencies arose recently in both the human and chimpanzee lineages, highlighting the importance of experimentally defining which cellular and molecular differences are human derived to inform our understanding of human evolution.

## Discussion

Loss-of-function screens have provided deep insights into the genes that regulate the development of model organisms. Here, we extended the scope of genetic screens to human and chimpanzee PSCs and examined whether the requirements for essential genes could differ in closely related species. By performing paired genome-wide CRISPRi screens, we uncovered a divergent landscape of genetic dependencies. Despite human and chimpanzee PSCs being similar in their cellular morphology, response to *in vitro* differentiation protocols, and core essential-ome, we identified 75 genes with divergent roles in controlling cellular proliferation. We observed that many of these genes were organized in coherent protein complexes and biochemical pathways, a metric of internal consistency that demonstrated the technical quality of our screens. Our data thus comprise a rich resource that interfaces with existing studies of gene regulation and chromatin states and provides a functional genomics guide for future candidate gene approaches.

Our work further demonstrates the capacity for comparative screening approaches to reveal latent molecular differences between closely-related species. The keys to the success of our approach include: 1) the scalability of CRISPRi sgRNA libraries, which target each gene with multiple unique sgRNAs, enabling the comprehensive and unbiased exploration of species-specific differences in genetic dependencies 2) the ability to design sgRNAs that target conserved genomic sequences in human and chimpanzee cells, and 3) the use of cells lines from multiple individuals of each species to distinguish consistent species differences from individual and technical sources of variation. We found that human and chimpanzee PSCs exhibit differential sensitivity to depletion of key components of cell cycle, lysosomal function, UFMylation, and histone modification. We further applied our comparative screening approach to orangutans, an outgroup to both humans and chimpanzees. By performing a three species comparison, we inferred the evolutionary history of several divergent species-specific molecular phenotypes. We found that robustness to depletion of *CDK2, CDK4,* and *Cyclin E1* and sensitivity to depletion of *ATP6AP1* were unique to humans, and thus likely arose after the last common ancestor of humans and chimpanzees. In contrast, depletion of *KAT6A* and *UFL1* led to changes in growth that were specific to chimpanzees. Thus, our ability to distinguish which species-differences arose within the human lineage is an indicator of their potential relevance to human phenotypic evolution.

How might the genetic dependencies we observed in PSCs relate to organismal differences that manifest during development? Intriguingly, we found that human-specific robustness to depletion of cell cycle factors persisted in neural progenitor cells. The G1-phase length hypothesis proposes that factors which lengthen G1 duration in NPCs increase the probability of differentiation towards non-proliferative neuronal fates, while factors reducing G1 length promote proliferative self-renewal of NPCs (Calegari et al., 2005; Dehay and Kennedy, 2007; Taverna et al., 2014). Indeed, loss of CDK2 or CDK4 in mouse NPCs prolongs G1 length and causes premature neuronal differentiation at the expense of self-renewal (Lim and Kaldis, 2012). Conversely, exogenous overexpression of *CDK4* and *Cyclin D1* in mouse and ferret reduces G1 length, promotes self-renewing divisions in basal progenitor cells, and results in increased brain size and cortical area, while preserving a structurally normal, six-layered cortex (Nonaka-Kinoshita et al., 2013). These studies underscore the influence of inputs to the G1/S transition on brain expansion during development. However, whether this developmental mechanism changed specifically in recent human evolution remained unexplored. Our demonstration that human NPCs are more likely than chimpanzee NPCs to continue cycling upon equivalent repression of *CDK2* and *Cyclin E1* connects proposed developmental mechanisms to molecular changes that occurred in human evolution. Our results, in concordance with previous studies, support a general framework where a combination of intrinsic developmental tempo and external environmental inputs govern the length of G1-phase in neural progenitor cells (Figure 7A). Factors promoting rapid G1/S transition, such as mitogen signaling and cyclin-CDK activity, will promote neural progenitor self-renewal and lead to an expansion of the progenitor field. Similarly, environmental stressors that lengthen G1-phase such as nutrient limitation and DNA damage can trigger neural progenitor differentiation. Molecular changes that make human NPCs less sensitive to these differentiation-promoting signals, as we observed via genetic perturbations, could bias human NPCs to increased proliferation in more complex *in vivo* environments. Future studies can thus assess the response of human and chimpanzee NPCs to a wider range of genetic and physiological perturbations to provide further insights into the evolutionary mechanisms by which the proliferative capacity of NPCs has increased along the human lineage.

**Figure 7.**
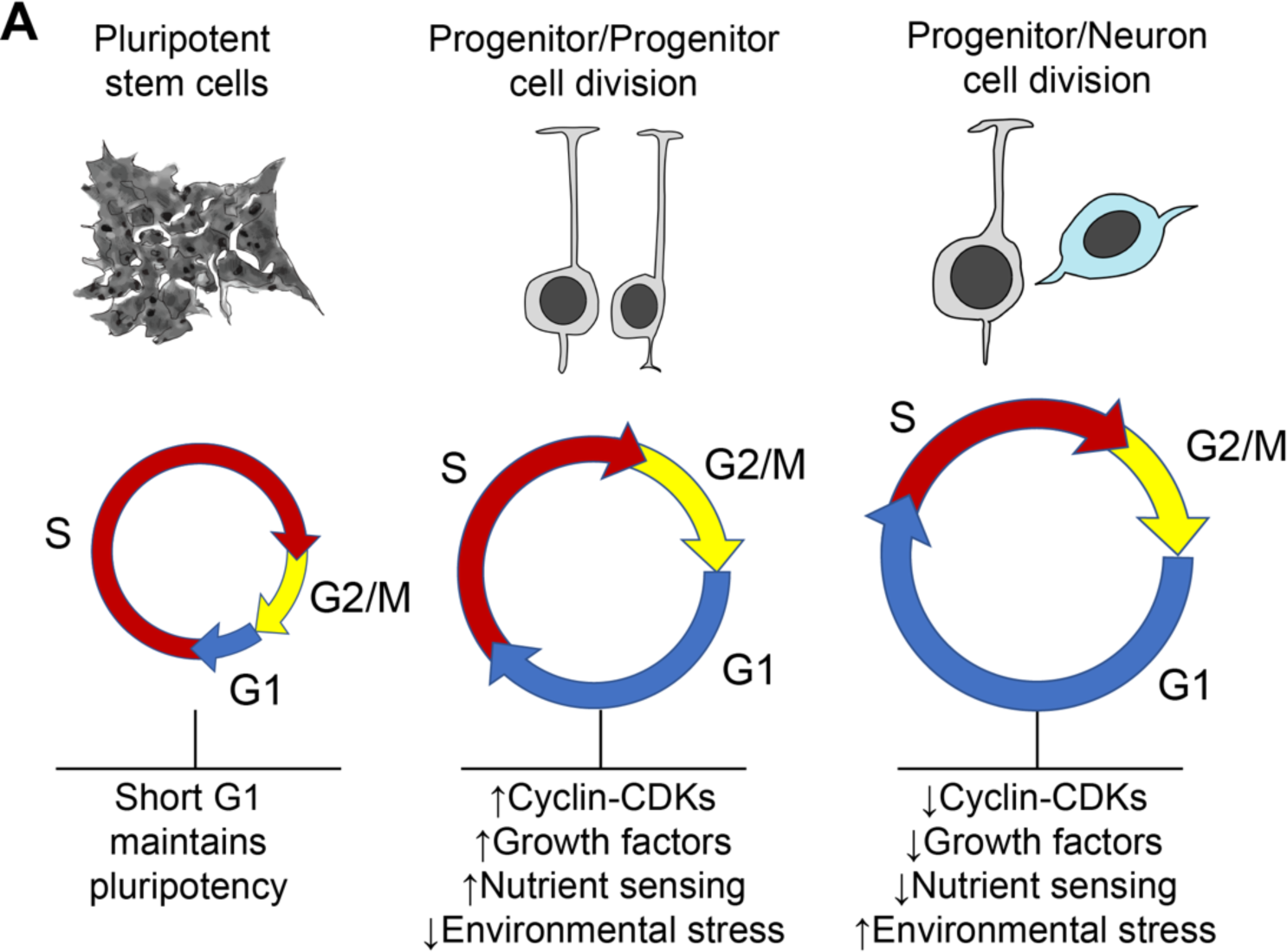
Model for inputs to neural progenitor differentiation. (A) Schematic for changes in cell cycle length and proportion in pluripotent stem cells and neural progenitor cells with intrinsic and extrinsic inputs that can potentially influence the duration of G1 phase.

The endeavor to study the molecular basis of human evolution has been compared to searching for needles in a haystack, as human-specific genetic variants and gene expression changes are numerous and predominantly neutral (Varki and Altheide, 2005). By contrast, our finding that human and chimpanzee PSCs exhibit distinct genetic dependencies, even for genes that lack clear expression or protein-coding sequence divergence, provides a complementary approach for isolating recently-evolved functional changes in human gene networks. This conceptual advance mirrors the progression of cancer genetics research from sequencing and transcriptomics efforts such as TCGA (Weinstein et al., 2013) to functional genetics-based efforts such as DepMap (Pacini et al., 2021; Tsherniak et al., 2017). Moreover, while driver mutations can be identified in tumors based on their independent recurrence, human evolution has occurred only once, highlighting the added value of a functional genomics platform. Our approach can be readily applied in differentiated cell types and interfaced with higher dimensional measurements of cell phenotypes (Adamson et al., 2016; Dixit et al., 2016; Feldman et al., 2019), opening the door to future efforts for understanding molecular control of species differences across stages of development.<colcnt=2>

**STAR METHODS.** KEY RESOURCES TABLE.

KEY RESOURCES TABLE
| REAGENT or RESOURCE | Source | Identifier |
| --- | --- | --- |
| **Antibodies** |  |  |
| anti-pS6, rabbit, 1:1,000 | Cell Signaling | Cat# 2211S; RRID: AB_331679 |
| anti-GAPDH, rabbit, 1:1,000 | Proteintech | Cat# 10494-1-AP; RRID: AB_2263076 |
| anti-PAX6, human, 1:50 | Miltenyi | Cat# 130-123-328; RRID: AB_2819477 |
| anti-B2M, mouse, 1:20 | BioLegend | Cat# 316312; RRID: AB_10641281 |
| anti-Nestin, mouse, 1:20 | BioLegend | Cat# 656805; RRID: AB_2566381 |
| Goat anti-Rabbit IGG secondary IRDye 680RD, 1:15,000 | LI-COR | Cat# 926-68071; RRID: AB_10956166 |
| Goat anti-Rabbit IGG secondary IRDye 800CW, 1:15,000 | LI-COR | Cat# 926-32211; RRID: AB_621843 |
| **Bacterial and virus strains** |  |  |
| MegaX Competent Cells | ThermoFisher | Cat# C640003 |
| Stellar Competent Cells | Takara | Cat# 636766 |
| **Chemicals, peptides, and recombinant proteins** |  |  |
| TransIT-LT1 Transfection Reagent | Mirus Bio | Cat# MIR2300 |
| Lipofectamine Stem Transfection Reagent | ThermoFisher | Cat# STEM00008 |
| SB431542 | Miltenyi | Cat# 130-106-543 |
| Noggin | Miltenyi | Cat# 130-103-456 |
| ROCK inhibitor thiazovivin | Stem Cell Tech. | 72252 |
| Puromycin | Goldbio | P-600-100 |
| Chroman 1 | Tocris | Cat# 7163 |
| Emricasan | Selleck Chemicals | Cat# S7775 |
| Polyamine Supplement | Sigma | Cat# P8483 |
| Trans-ISRIB | Tocris | Cat# 5284 |
| **Critical commercial assays** |  |  |
| Macherey-Nagel NucleoSpin Blood XL | ThermoFisher | Cat# NC1105387 |
| Superscript IV VILO | ThermoFisher | Cat# 11756050 |
| DyNAmo ColorFlash SYBR Green | ThermoFisher | Cat# F416L |
| Click-iT Plus EdU Alexa Fluor 647 Flow Cytometry Assay | ThermoFisher | Cat# C10635 |
| BD Cytotfix/Cytoperm | ThermoFisher | Cat# BDB554714 |
| **Deposited Data** |  |  |
| Raw sequencing data from bulk RNA-seq | This paper | GEO |
| Processed sequencing data from bulk RNA-seq | This paper | Table S2 |
| **Experimental Models: Cell lines** |  |  |
| Chimpanzee iPSC 3624K dCas9-KRAB-mCherry | This paper | N/A |
| Chimpanzee iPSC 3624K dCas9-KRAB-BFP | This paper | N/A |
| Chimpanzee iPSC 3649 dCas9-KRAB-BFP | This paper | N/A |
| Chimpanzee iPSC 40290F dCas9-KRAB-mCherry | This paper | N/A |
| Chimpanzee iPSC 8861G dCas9-KRAB-mCherry | This paper | N/A |
| Chimpanzee iPSC 8861G dCas9-KRAB-BFP | This paper | N/A |
| Chimpanzee iPSC Pt-5c dCas9-KRAB-BFP | This paper | N/A |
| Human iPSC 20961B dCas9-KRAB-BFP | This paper | N/A |
| Human iPSC 20961B dCas9-KRAB-GFP | This paper | N/A |
| Human iPSC 21792A dCas9-KRAB-BFP | This paper | N/A |
| Human iPSC 23555A dCas9-KRAB-BFP | This paper | N/A |
| Human iPSC 23555A dCas9-KRAB-GFP | This paper | N/A |
| Human iPSC 28128B dCas9-KRAB-BFP | This paper | N/A |
| Human iPSC 28128B dCas9-KRAB-GFP | This paper | N/A |
| Human ESC H1 dCas9-KRAB-BFP | This paper | N/A |
| Human iPSC WTC11 dCas9-KRAB-BFP | This paper | N/A |
| **Oligonucleotides** |  |  |
| See Table S6 for oligonucleotides used in this study | This paper | N/A |
| **Recombinant DNA** |  |  |
| pX458 | Addgene | Cat# 48138 |
| Gen3-AAVS1 dCas9-XTEN-KRAB-P2A-BFP | Addgene | Cat# 127968 |
| pEF1-BCL-XL | (Li et al., 2018) |  |
| pCRISPRia-v2 | Addgene<br>(Adamson et al., 2016) | Cat# #84832 |
| **Software and Algorithms** |  |  |
| CRISPRi screen processing for sgRNA counts | (Horlbeck et al., 2016) | https://github.com/mhorlbeck/ScreenProcessing |
| MaGeCK | (Li et al., 2014) | https://sourceforge.net/projects/mageck/ |
| FlashFry | (McKenna and Shendure, 2018) | https://github.com/mckennalab/FlashFry |
| Cutadapt | (Martin, 2011) | https://cutadapt.readthedocs.io/en/stable/ |
| kallisto | (Bray et al., 2016) | https://pachterlab.github.io/kallisto/ |
| gffread | (Pertea and Pertea, 2020) | https://github.com/gpertea/gffread |
| tximport | (Soneson et al., 2015) | https://bioconductor.org/packages/release/bioc/html/tximport.html |
| DESeq2 | (Love et al., 2014) | https://bioconductor.org/packages/release/bioc/html/DESeq2.html |

## CONTACT FOR REAGENT AND RESOURCE SHARING

Further information and requests for resources and reagents should be directed to and will be fulfilled by the Lead Contact, Alex A Pollen.

### Cell Lines

All cell lines used in this study are listed in Supplementary Table 1.

### Primers

All primers used in this study are listed in Supplementary Table 6.

### Media Formulations

mTESR1 was purchased from Stem Cell Technologies (cat. 85850) and supplemented with 100 units/ml penicillin, 100 µg/ml streptomycin, and 292 µg/ml L-glutamine (Gibco, cat. 10378016). StemFlex was purchased from Gibco (cat. A3349401) and supplemented with 100 units/ml penicillin, 100 µg/ml streptomycin, and 292 µg/ml L-glutamine. HEK293Ts were cultured in DMEM (ThermoFisher, cat. 11965118) supplemented with 10% FBS (VWR, cat. 97068-085, lot 043K20), 100 units/ml penicillin, 100 µg/ml streptomycin, and 292 µg/ml L-glutamine. Neuronal differentiation media was prepared as described in (Nolbrant et al., 2017), with DMEM F/12 (ThermoFisher, cat. 21331020), CTS Neurobasal Medium (ThermoFisher, cat. A1371201), 1x N-2 supplement CTS (ThermoFisher, cat. A1370701), 10 µM SB431542 (StemMACS, TGFβ inhibitor; Miltenyi, cat. 130-106-543), 100 ng/ml Noggin (recombinant human; Miltenyi, cat. 130-103-456), and 100 units/ml penicillin, 100 µg/ml streptomycin, and 292 µg/ml L-glutamine. Thiazovivin (Stem Cell Technologies, cat. 72252) was included at a concentration of 2 μM during passaging. For single cell sorting and lipofection, we used the CEPT cocktail (Chen et al., 2021) consisting of 50 nM chroman 1 (Tocris Bioscience, cat. 7163), 5 µM emricasan (Selleck Chemicals, cat. S7775), polyamine supplement diluted 1:1,000 (Sigma-Aldrich, cat. P8483), and 0.7 µM trans-ISRIB (Tocris Bioscience, cat. 5284).

### Construction of CRISPRi cell lines

All wildtype cell lines tested negative for mycoplasma prior to the start of cell line engineering. The CRISPRi effector protein dCas9-KRAB was introduced into either the CLYBL or AAVS1 safe harbor locus (Cheung et al., 1980; Hockemeyer et al., 2009; Philpott et al., 2002; Smith et al., 2008) via lipofection of three plasmids: 1) A modified version of pX458 (Addgene #48138), containing both Cas9 nuclease and a sgRNA targeting either CLYBL or AAVS1. The sgRNA spacer was modified from the original plasmid by cutting with type IIS restriction endonuclease BbsI-HF. Complementary oligos containing the sgRNA spacer and proper overhangs were annealed and ligated with T4 ligase. 2) A modified version of the Gen3-AAVS1 vector from (Mandegar et al., 2016) containing dCas9-XTEN-KRAB-P2A-BFP driven by the chicken beta-actin (CAG) promoter, flanked by homology arms to CLYBL (Addgene #127968) or AAVS1. 3) pEF1-BCL-XL, a plasmid expressing BCL-XL, the anti-apoptotic isoform of BCL2L1, from the EF-1a promoter. Lipofection was performed as follows: two days prior to transfection, cells were switched into mTESR1 media on a non-passaging day. One day prior to transfection, ∼400,000 cells were plated into a Matrigel-coated (Corning, cat. 354230) 6-well plate with mTESR1 supplemented with 2 μM thiazovivin. On the day of transfection, a 3 μg mixture of plasmids 1-3 was made at a mass ratio of 5:5:1, added to a mixture of 96 μl Opti-MEM (Gibco, cat. 31985062) and 4 μl Lipofectamine Stem (ThermoFisher, cat. STEM00003), and incubated at room temperature for 10 minutes. Media was aspirated from the PSC plate and replaced with 2 ml Opti-MEM supplemented with CEPT. The lipid/plasmid DNA complexes were then added to plate and incubated for 4 hours, after which 2 ml of mTESR1 supplemented with CEPT was overlaid. 24 hr post-transfection, media was replaced with StemFlex supplemented with CEPT. 48 hr post-transfection, media was replaced with StemFlex supplemented with 2 μM thiazovivin, and cells were then passaged for 10-14 days to dilute out the transfected plasmids. Single cell clones and one polyclonal population per cell line were then sorted on a Sony MA900 (see Table S1). Expanded populations were cryopreserved in Bambanker preservation media (ThermoFisher, cat. 50999554) and functionally validated with an sgRNA targeting B2M, a non-essential surface marker. B2M levels were measured by staining with an APC anti-human β2-microglobulin antibody (BioLegend, cat. 316312).

### Lentivirus production, concentration, and titration

Lentivirus for CRISPRi screening was produced in HEK293T cells. HEK293Ts were seeded at a density of 80,000 cells/cm^2^ 24 hr. prior to transfection in 15 cm dishes. Next, each dish was transduced with 20 μg sgRNA library, 6.75 μg standard lentivirus packaging vectors, and 81 μl Mirus transfection reagent (VWR, cat. 10767-122) in Opti-MEM. 24 hr. post-transfection, media was replaced and supplemented with 1X ViralBoost (Alstem, cat. VB100). Supernatant was collected at 48 hr. post-transfection and concentrated 1:10 with Lenti-X Concentrator (Takara Bio, cat. 631231). Concentrated lentivirus was titered in PSCs based on BFP expression 3 days post-infection using a flow cytometer.

### Pooled genome-wide CRISPRi screening

CRISPRi PSCs expressing dCas9-KRAB were dissociated with Accutase (Innovative Cell Technologies, cat. AT104-500), resuspended in StemFlex supplemented with 2 μM thiazovivin and 5 μg/ml polybrene (Sigma-Aldrich, cat. TR-1003-G), transduced with the lentiviral hCRISPRi-v2 sgRNA library at a target infection rate of 25-40%, and plated in Matrigel-coated 5-layer cell culture flasks (Corning, cat. 353144) at a density of 65,000-80,000 cells/cm^2^. The following day, StemFlex medium was replaced. Two days after infection, cells were dissociated with Accutase, resuspended in StemFlex supplemented with 2 μM thiazovivin and 1.5 μg/ml puromycin (Goldbio, cat. P-600-100), and plated in 5-layer cell culture flasks. The following day, medium was replaced with StemFlex supplemented with 1.5 μg/ml puromycin. Four days after infection, 100 M cells were harvested for the initial time point (t0), while 250-300 M cells were resuspended in StemFlex supplemented with 1.5 μg/ml puromycin and plated in 5-layer cell culture flasks (>1000x sgRNA library representation). Selection efficiency was assessed by flow cytometry (>70% BFP+). Every two days, cells were dissociated with Accutase, resuspended in StemFlex supplemented with 2 μM thiazovivin, and plated at a density of 80,000-100,000 cells/cm^2^. Technical replicates were cultured separately for the duration of the screen. After 10 days of growth, 150 M cells from each technical replicate were harvested for the final time point (tfinal). Genomic DNA was isolated from frozen cell pellets using the Macherey-Nagel NucleoSpin Blood XL kit (Macherey-Nagel). Isolated DNA was quantified using a NanoDrop (ThermoFisher) and the sgRNA expression cassette was amplified by 22 cycles of PCR using NEBNext Ultra II Q5 Master Mix (NEB) and primers containing Illumina P5/P7 termini and sample-specific TruSeq indices. Each sample was distributed into 150-200 individual 100 µl reactions in 96-well plates, each with 10 μg genomic DNA as input. Following amplification, reactions from each sample were pooled and a 100 μl aliquot was purified using AMPure XP beads (Beckman-Coulter) with a two-sided size selection. Purified libraries were quantified using a Qubit (ThermoFisher) and sequenced using an Illumina HiSeq 4000 (SE50, 5% PhiX) with a custom sequencing primer (oCRISPRi_seq V5).

### Data analysis for pooled genome-wide CRISPRi screens

Sequencing data were aligned to hCRISPRi-v2 and quantified using the ScreenProcessing pipeline (https://github.com/mhorlbeck/ScreenProcessing) (Horlbeck et al., 2016). sgRNA counts were then processed using MAGeCK (Li et al., 2014) test (-norm-method control-remove-zero control-gene-test-fdr-threshold 0.10 −remove-zero-threshold 50 −gene-lfc-method alphamean) and separately using a custom analysis method inspired by MAGeCK. Briefly, sgRNA counts were normalized by the median ratio method. Mean-variance modeling was performed with non-targeting sgRNAs as the control group, and sample mean and variance values were used to parameterize a negative binomial distribution. *P*-values were then calculated for each sgRNA based on the tail probability of the negative binomial and *P*-value cut-offs were chosen such that 95% of non-targeting sgRNAs were not significant.

For each gene, sgRNAs were filtered by two criteria: 1) perfect alignment to both the human (hg38) and chimpanzee (panTro6) reference genomes, as determined by FlashFry (McKenna and Shendure, 2018), and 2) significance according to the negative binomial distribution. Only sgRNAs passing both filters were retained for analysis, resulting in a variable number of sgRNAs per gene (0-5 sgRNAs). The remaining sgRNA counts were converted to log2 fold-change and averaged to produce a gene score. Significance testing for gene scores was performed by bootstrapping non-targeting sgRNAs, with groups of 1-5 random non-targeting sgRNAs assigned to each control gene to select candidates from the initial genome-wide screens.

Essential genes for each screen (Fig. 1B) were determined by the degree of depletion among sgRNAs targeting the gene. To facilitate equal comparison among screens, the top 3000 most depleted genes in each screen were defined as essential (with mean sgRNA depletion greater than 4-fold for all such genes). In addition, genes with mean sgRNA depletion less than 2-fold were defined as non-essential. Each intersection set was then constructed based on two inclusion criteria: genes were required to be essential for each member of the set and non-essential for all non-members of the set.

For comparison of technical replicates (Fig. 1B), identification of shared essential genes (Fig. 1B), and assessment of screen performance using DepMap Public 21Q4 gene sets (Fig. 1C), all hCRISPRi-v2 sgRNAs, including those with mismatched targets in the panTro6 reference genome, were included for analysis. Mismatches specific to each individual cell line were not considered, as high-quality genome sequences of each cell line were not available. However, these mismatches should in theory affect only a small fraction of all sgRNAs. For identification of candidate genes with species-specific effects on proliferation (Fig. 1D), only sgRNAs with perfect-match targets in the panTro6 reference genome (77.4%, *n* = 79417/102640) and transcriptional start sites targeted by at least three sgRNAs after excluding mismatched sgRNAs (86.7%, *n* = 17804/20528), were retained for analysis.

For screen analysis using MAGeCK, genes with false discovery rates (FDRs) less than 10% for both individuals within a species and FDRs greater than 25% in both individuals from the opposite species were considered as candidates for validation screening (*n* = 418 genes).

### CEV-v1 validation screening library design

To validate genome-wide screens, a Comparative Essential Validation (CEV-v1) sgRNA library consisting of 9,692 sgRNAs targeting 963 candidate species-specific essential or proliferation suppressor genes was constructed. hCRISPRi-v2 sgRNAs with perfect-match targets in panTro6 exhibiting significant depletion or enrichment in the genome-wide screens were retained in CEV-v1 (*n* = 3589 sgRNAs). In addition, new sgRNAs with perfect-match target sites in the human (hg38) and chimpanzee (panTro6) reference genomes were chosen based on their position relative to the FANTOM-annotated transcriptional start site (Horlbeck et al., 2016) on-target activity predicted by DeepHF (Wang et al., 2019), and off-target potential predicted by a genome-wide search of mismatched target sites (Jost et al., 2020; McKenna and Shendure, 2018) in both reference genomes. Briefly, after performing off-target filtering (one perfect-match target, CRISPRi specificity score > 0.20, maximal predicted activity at any off-target site < 0.80), candidate sgRNAs were categorized by their position relative to the FANTOM TSS and then ranked by their DeepHF score. A threshold DeepHF score was imposed by excluding sgRNAs with predicted activities less than one standard deviation below the mean of all candidate sgRNAs (minimum score: 0.4378). Eight sgRNAs were selected for each gene as well as 1845 non-targeting sgRNAs from hCRISPRi-v2. Oligonucleotide pools were designed with flanking PCR and restriction sites (BstXI, BlpI), synthesized by Agilent Technologies, and cloned into the sgRNA expression vector pCRISPRia-v2 (Addgene #84832) as described previously (Gilbert et al., 2014).

### Pooled validation CRISPRi screening

Validation screens were performed in conditions consistent with the genome-wide screens. Briefly, CRISPRi PSCs expressing dCas9-KRAB were dissociated with Accutase, resuspended in StemFlex supplemented with 2 μM thiazovivin and 5 μg/ml polybrene, transduced with the lentiviral CEV-v1 sgRNA library at a target infection rate of 25-40%, and plated in Matrigel-coated 3-layer cell culture flasks (Corning, cat. 353143) at a density of 65,000-80,000 cells/cm^2^. Cells were dissociated, plated, selected with puromycin, and grown on the same schedule as used for the genome-wide screens. Technical replicates were cultured separately for the duration of the screen and >1000x sgRNA library representation was maintained. For individuals screened twice (H20961B, H23555A, H28126B, C3624K, C8861G), replicate screens were performed independently at the Whitehead Institute and UCSF.

### Data analysis for CEV-v1 validation screens

Sequencing data were aligned to CEV-v1 and quantified using the ScreenProcessing pipeline and MAGeCK. A matrix containing sgRNA counts from all CEV-v1 screens (excluding C3649 and Pt5-C due to non-responsiveness to *MDM2*/*p53* perturbations) was assembled and used as input for differential analysis by DESeq2. Briefly, each sample was annotated by species, individual, and timepoint and a design matrix was created to model the species-specific effect of time (t0 vs. tfinal) while controlling for individual effects (modeled as fixed effects) within each species. The human and chimpanzee species terms were then contrasted to extract a Benjamini–Hochberg-adjusted *P-*value and log2 fold-change for each sgRNA. sgRNA adjusted *P-*values were combined into gene FDRs using alpha-robust rank aggregation (α-RRA) from MAGeCK and the α threshold (to remove the effect of insignificant sgRNAs from the assessment of gene significance) was set according to the fraction of sgRNAs with an adjusted *P-*value < 0.01. For each gene, log2 fold-change was computed as the mean of the four sgRNAs with the largest absolute fold-change. To exclude genes with shared effects from being erroneously called as species-specific, any gene with an FDR in both the human and chimpanzee species terms less than the highest FDR for any gene with at least one sgRNA passing the α threshold in α-RRA was discarded. For each gene in the set of 75 genes with species-specific effects reported in Fig. 2, we required that three conditions be met in the chimpanzee–human contrast term: (1) gene FDR < 0.01, (2) at least three sgRNAs targeting the gene pass the α threshold in α-RRA, and (3) gene log2 fold-change difference ≥ 0.75 between species. We used the STRING database v11.5 (Szklarczyk et al., 2021) to identify known and predicted protein–protein interactions among this set of 75 genes.

To quantify sources of variation in CEV-v1 screens, a matrix of sgRNA counts was assembled as described above and normalized using edgeR (Robinson et al., 2010) calcNormFactors. Normalized sgRNA counts were then prepared for linear modeling using variancePartition voomWithDreamWeights and a linear mixed model was fit using variancePartition fitExtractVarPartModel. The categorical variables species, individual, and timepoint were modeled as random effects. For each gene (Fig. S2B), gene-level estimates of variance were determined by computing the mean variance attributable to each variable for all sgRNAs targeting that gene.

### Quantitative RT-PCR

Biological triplicates were grown in a 6-well plate and infected with sgRNAs (see Table S2) at an MOI of ∼0.3. 48h post-infection, cells were expanded in 2 μg/ml puromycin and allowed to recover for 48 hours. At day 5 post-infection, sgRNA+ cells were sorted to purity based on BFP+ expression on a Sony MA900 cell sorter. Cells were then allowed to recover for 48-96 hours, until they reached ∼60-80% confluence on a 6-well plate. RNA was extracted with a Direct-zol RNA miniprep kit (Zymo Research, cat. R2051). RNA was reverse transcribed with SuperScript IV VILO (ThermoFisher, cat. 11756050), and cDNA was amplified with the DyNAmo ColorFlash SYBR Green kit (ThermoFisher, cat. F416L). Primers for GAPDH were used as loading controls and no-RT controls were performed to control for genomic DNA contamination. Amplifications were performed in duplicate and quantified on a QuantStudio Flex 7 Real-Time PCR system in 96-well plates.

### Copy number variation analysis

Chromosomal copy number variations (CNV) were inferred with the InferCNV R package (version 1.2.1), which predicts CNVs based on gene expression data. InferCNV was run in ‘subclusters’ analysis mode using ‘random_trees’ as the subclustering method with gene expression quantified for both species by alignment to the hg38 reference genome. Average gene expression across all six individuals in each species was used as the background column. The cut-off for the minimum average read count per gene among reference cells was set to 1, per software recommendation for bulk RNA-seq data. CNV prediction was performed with the ‘i6’ Hidden Markov Model, whose output CNV states were filtered with the included Bayesian mixture model with a threshold of 0.1 to find the most confident CNVs. All other options were set to their default values.

To check for copy number variation at a selected set of cell cycle-related genes, we analyzed whole-genome shotgun sequencing data. Genomic DNA from all libraries sequenced on an Illumina sequencer in 151 bp paired-end mode was provided for analysis courtesy of the laboratory of Gregory Wray and mapped to chimpanzee reference panTro6 (Kronenberg et al., 2018) using bwa-mem2 (https://ieeexplore.ieee.org/document/8820962) with default parameters. PicardTools (https://broadinstitute.github.io/picard) was used to add read group information and mark duplicates, and baseline coverage histograms were generated using BEDTools genomecov (Quinlan and Hall, 2010), from which the 5th, 50th, and 95th percentile of coverage for each library, both genome-wide and across chromosome X, were extracted. Gene-level features for all genes listed as cyclins, cyclin dependent kinases, and class III Cys-based CDC25 phosphatases in the HGNC database (Tweedie et al., 2021) were selected from a recent chimpanzee gene annotation (Mao et al., 2021) and the coverage at each base across the full length of each gene in the set for each library was counted and summed using samtools mpileup (Li et al., 2009). For this step, only primary alignments containing mapped reads not marked as duplicates, with minimum map quality of 20, were considered (samtools view −F1284 −q20).

### RNA-seq library prep and analysis

Human (WTC11) and chimpanzee (C3649) cells were infected with individual sgRNAs targeting KAT6A or BRPF1. Cells were expanded for 5 days. At day 5, ∼1M sgRNA expressing cells were sorted on a Sony MA900 cell sorter based on BFP+ expression. Cells were then pelleted, snap-frozen and stored at −80°C. High quality RNA was extracted by adding RNAse-free Trizol (ThermoFisher, cat. 15596026) to each pellet and processing with the Zymo Research Direct-zol RNA miniprep kit (Zymo Research, cat. R2050). RNA-seq was performed using the Illumina TruSeq Stranded Total RNA kit (Illumina, cat. 20020599) according to the manufacturer’s instructions, with the exception of the final PCR step for which only 10 cycles were used to prevent overamplification. The final pooled library was sequenced with 50 bp single end reads on a HiSeq 2500.

Raw bulk RNA-seq reads from knockdown experiments and wild-type chimpanzee and human iPSCs were adapter-trimmed using cutadapt (Martin, 2011) (with option −b AGATCGGAAGAGCACACGTCTGAACTCCAGTCA) and then pseudo-aligned to species-specific transcriptomes using kallisto (Bray et al., 2016) with options −single −l 200 −s 20. Transcriptomes were extracted from species-specific gtf annotations using the gffread utility (Pertea and Pertea, 2020) using the −w output option. Human transcripts were obtained from the Gencode (Frankish et al., 2021) comprehensive gene annotation v36 (GTF), using genome assembly hg38 (Schneider et al., 2017), and the chimpanzee annotation was obtained from a recent study that produced a hierarchical alignment of primate genome assemblies (Mao et al., 2021) and annotated the assemblies using the Comparative Annotation Toolkit (Fiddes et al., 2018).

To ensure consistency of gene names across the annotations, we downloaded the set of gene aliases from the HUGO Gene Nomenclature Committee website (www.genenames.org; (Tweedie et al., 2021) and searched for gene names present in the chimpanzee but missing in the human annotation, mapped to aliases present in the human annotation. This led us to rename five genes in the non-human primate annotations (DEC1 to BHLHE40, DUSP27 to DUSP29, AC073585.2 to FAM24B, LOR to LOXL2, and TNRC6C-AS1 to TMC6); we also renamed CCNP in the human annotation to CNTD2.

After counting transcript abundances using kallisto, we converted them to gene counts using the tximport command in the tximport R package (Soneson et al., 2015) with the options type=’kallisto’ and countsFromAbundance=’no’. We then created one human and one chimpanzee data set in DESeq2 (Love et al., 2014) contrasting gene knockdowns with wild-type gene expression in two replicates of the same cell line in each. We extracted VST-transformed counts for plotting using the function vst with option blind=TRUE and ran the DESeq linear model fitting using the function DESeq with betaPrior=TRUE.

GO enrichment was performed on the set of genes overexpressed in cells expressing sgRNAs targeting *KAT6A* or *BRPF1*. Overexpressed genes were defined as genes with fold-change > 2 in sgRNA expressing cells. Fold-change values were averaged between cells expressing sgBRPF1 and sgKAT6A. GO enrichment analysis was performed with Enrichr (Chen et al., 2013) and significant terms from the MSigDB Hallmark gene set were most significant and relevant for chimpanzee overexpressed genes. Significant terms from WikiPathway Human and GO Molecular Function were used for human overexpressed genes, though TGF-beta signaling was also an enriched GO term in MSigDB, and genes in the cell differentiation category were partially overlapping with TGF-beta signaling genes. Morphogen activity genes were *CER1* and *NODAL*.

Gene expression changes were quantified for a set of 149 developmental marker genes with evidence from prior literature (Maguire et al., 2013). Among these genes, 21 were overexpressed by > 0.5 by log2 fold-change in either human or chimpanzee cells. 8/21 were further filtered out based on DESeq2 adjusted *P*-value. Fold-change values were averaged between cells expressing sgBRPF1 and sgKAT6A. Markers were clustered based on overexpression in both human and chimp or human alone. Markers were then clustered and colored based on their association with ectoderm, endoderm, mesoderm, or mixed lineages. Lastly, canonical pluripotency markers *OCT4*, *SOX2*, and *NANOG* were added for visual comparison, as none of these genes were significantly upregulated.

### Analysis of protein-coding and gene expression changes

To obtain coding sequences for homologous transcripts from human and chimpanzee reference genomes, we downloaded human protein and transcript sequences from Gencode release 36 (Frankish et al., 2021) and chimpanzee protein and transcript sequences from the Comparative Annotation Toolkit (Fiddes et al., 2018) annotation on reference version panTro6 produced as part of a recent study (Mao et al., 2021). For each human transcript of each protein coding gene, we obtained the transcript sequence and its canonical translation, and we then extracted the corresponding chimpanzee transcript and canonical translation by matching the Ensembl transcript ID to its chimpanzee counterpart (“source transcript” field in the chimpanzee gene annotation). For both the human and chimpanzee sequence of each transcript, we then compared the translated sequence at every possible start codon and frame to the canonical amino acid sequence, determining the start codon and frame that produced the canonical amino acid sequence to be “correct” and removing bases thus found to belong to the 5’ or 3’ UTR (upstream of the correct start codon or downstream of the correct stop codon).

With coding sequences for homologous transcripts, we then aligned the human and chimpanzee protein sequences using the pairwise2 module from Biopython (Cock et al., 2009) with a BLOSUM62 substitution matrix. We then deleted codons in transcripts corresponding to amino acids that aligned to a gap in the other amino acid sequence. Finally, we deleted stop codons from the ends of sequences. We then wrote out each pairwise alignment to a control file for PAML (Yang, 2007) and calculated relevant statistics, including dN, dS, N, and S, using PAML’s implementation of the Yang and Nielson 2000 (yn00) algorithm (Yang and Nielsen, 2000). Finally, to avoid undefined values, we set dS to 1/S where dS was zero and selected the median dN value and median dN/dS value per gene for analysis.

Distributions of dN and dN/dS were compared for the full set of genes, DepMap common essential genes, and validation screen hits by two-sided Kolmogorov-Smirnov test (ks.test in R).

### Western Blot

Human (28128B) and chimpanzee (40280L) PSCs were infected with lentiviral constructs containing sgRNAs targeting ATP6AP1 or ATP6AP2. 48 hours post-infection, cells expressing sgRNA were selected for via addition of 1.5 μg/ml puromycin. Cells were then recovered in normal growth media from 4 days post-infection to 6 days post-infection. Cells were harvested at 6 days post-infection, along with separate wells of three wild-type human (H1, 21792A, 28128B) and three wild-type chimpanzee (3624K, 40280L, 8861G) cell lines. Cells from each well of a 6-well plate were lysed in ∼250 μl ice-cold RIPA buffer + protease inhibitor (ThermoFisher, cat. A32965). After 30 minutes of incubation in lysis buffer at 4°C, cells were centrifuged at 16,000 x g for 5 minutes at 4°C. Supernatant was collected and snap frozen in liquid nitrogen and stored at −80°C.

Protein concentrations in each lysate were quantified using a Bradford BCA kit (ThermoFisher, cat. PI23227) Lysate was normalized to 1 μg/μl in RIPA buffer. 30 μl of lysate was added to 10 μl of NuPage Sample Buffer (4x), heated to 70°C on a PCR thermocycler, and loaded onto a Bolt 4-12% polyacrylamide gel (ThermoFisher, NW04122BOX). The gel was run for 45 minutes at 165V in MOPS buffer. Protein was then transferred onto a nitrocellulose membrane (BioRad, cat. 1704270) with a Bio-Rad Trans-Blot Turbo (BioRad, cat. 1704150). The membrane was blocked with Intercept (PBS) Blocking Buffer (LI-COR, cat. 927-90003) for 1 hour at RT. Membrane was incubated overnight at 4°C with anti-pS6 primary antibody at a 1:1,000 dilution. Membrane was washed 3x with TBST and incubated with secondary antibody at 1:15,000 dilution. Membrane was washed 3x with TBST and imaged on a LI-COR Odyssey CLX. Afterwards, antibodies were stripped from the membrane with NewBlot IR Stripping Buffer (LICOR, cat. 928-40028). Membrane was reblotted with anti-GAPDH antibody at 1:1,000 dilution and incubated overnight at 4°C. Membrane was washed 3x with TBST and incubated with secondary antibody at 1:15,000 dilution. Membrane was washed 3x with TBST and imaged on a LI-COR Odyssey CLX.

### Cell cycle EdU staining

Cell cycle phase measurements were performed with the Click-iT Plus EdU Alexa Fluor 647 Flow Cytometry Assay Kit (ThermoFisher, cat. C10635). 10 μM EdU was added to cell cultures for 1 hr, after which cells were harvested with Accutase. The cell pellet was washed once with once with 500 μl PBS supplemented with 1% BSA, pelleted again, and fixed for 15 minutes at RT, protected from light. Cells were washed and permeabilized for 15 minutes at RT. EdU detection was then accomplished via click chemistry of an Alexa Fluor 647 coupled to picolyl azide. After 1 wash, 10 μg/ml of Hoechst 33342 (ThermoFisher, cat. H3570) was added and incubated for 15 minutes. Cells were then directly analyzed by flow cytometry. Data were analyzed with custom Matlab scripts – after filtering for viable cells and doublets, G1, S, and G2/M gates were manually drawn and saved with the function impoly() for each sample. sgRNA+ populations were determined by GFP+ expression, and identical G1, S, and G2/M gates were used for sgRNA+ cells and sgRNA-cells within each sample.

### Cell cycle drug treatments

For Fig. 3F, human iPSC line H28128B was used and chimpanzee iPSC line C40280L was used. Cells were infected with an sgRNA targeting FAM122A (see Table S2), with 15-30% of cells infected. No puromycin selection was performed. At day 4 post-infection, cells were Accutase passaged into StemFlex supplemented with 2 μM thiazovivin and drug, with prexasertib (Chk1i) or adavosertib (Wee1i) added at 62 nM. At day 6, cells were replated and fresh drug was added to ensure removal of dead cells. At day 8, the fraction of sgRNA+ (BFP+) surviving cells were analyzed by flow cytometry.

For Fig. 3G, human iPSC line H21792A and chimpanzee iPSC C40280L were co-cultured with a 50/50 initial seeding density. After one normal passage, cells were Accutase passaged into StemFlex supplemented with 2 μM thiazovivin and drug, with prexasertib or adavosertib added at 125 nM. 24hr after drug treatment, cells were replated and fresh drug was added to ensure removal of dead cells. 48 hr post drug treatment, the ratio of human cells (GFP+) chimpanzee cells (mCherry+) was analyzed by flow cytometry.

### Neural progenitor cell differentiation

PSCs were differentiated into neural progenitor cells (NPCs) as described in (Nolbrant et al., 2017) with the following modifications. Differentiation media was made without ventral and caudalization patterning factors (sonic hedgehog agonist and GSK3i CHIR99021). PSCs were maintained on Matrigel (Corning, cat. 354230) prior to day 0 plating onto Lam-111 coated plates. Cells were seeded at 20,000 cells/cm^2^, as measured by Chemometec Nucleocounter NC-202, roughly twice the published density to ensure robust survival. NPCs were evaluated for purity at days 7-11 of differentiation with antibody staining against NPC markers Pax6 and Nestin. Cells were dissociated with Accutase, pelleted and washed, then fixed and permeabilized with the BD Cytofix/Cytoperm kit (ThermoFisher, cat. BDB554714). 100 μl cells were stained with 2 μl human anti-Pax6-APC (Miltenyi, cat. 130-123-328) + 5 μl mouse anti-Nestin-PE (Biolegend, cat. 656805) and evaluated by flow cytometry. In addition, NPCs were plated on μ-Slide 4 Well chambers (Ibidi cat. 80426), stained with antibodies against Pax6 and Nestin, and visualized by fluorescence microscopy on a RPI spinning disk confocal microscope.

### Orangutan CRISPRi growth comparison

CRISPRi machinery was engineered into orangutan PSCs (Field et al., 2019) at the AAVS1 locus via the three plasmid lipofection method described above (see Construction of CRISPRi cell lines). To account for mutations in the orangutan genome, the sgRNA for the Cas9 nuclease component was modified to perfectly match the orangutan AAVS1 locus. However, flanking regions were not modified. sgRNAs targeting *CDK2*, *CDK4*, *CCNE1*, *ATP6AP1*, *KAT6A*, and *UFL1* were transduced into human (28128B), chimpanzee (40280L), and orangutan PSCs via lentivirus at MOI ∼1. Cells were transduced in triplicate in 24-well plates and passaged every 2 days in ROCKi. At each passage, a portion of cells were quantified by flow cytometry on a BD LSRFortessa. The fraction of sgRNA+ expressing cells was determined based on the fraction of BFP+ cells. Measurements were collected until day 10 for *CDK2*, *CDK4*, *CCNE1*, and *ATP6AP1*. Measurements were collected until day 14 for *KAT6A* and *UFL1*. In parallel, cells transduced with sgRNAs targeting *CDK2*, *CDK4*, *CCNE1*, *ATP6AP1*, *KAT6A*, and *UFL1* and expanded into a 6 well plate. 48h post-infection, cells were expanded in 2 μg/ml puromycin and allowed to recover for 48 hours. At day 5 post-infection, sgRNA+ cells were sorted to purity based on BFP+ expression on a Sony MA900 cell sorter. RNA was extracted, cDNA was reverse transcribed, and qRT-PCR was used to quantify the degree of sgRNA-mediated depletion in biological triplicate (as described above in Quantitative RT-PCR).

## Data and code availability

Raw sequencing data are deposited on GEO accession number GSE212297.

All code for the analyses performed on the CRISPRi screens is publicly available at https://github.com/tdfair.

## Acknowledgments

We thank Eva Okrent for all illustrations contained in this manuscript. We thank Joseph Min for running the CaSpER CNV pipeline, Sofie Salama and Andrew Field for providing orangutan iPSCs, Gregory Wray for providing whole genome shotgun sequencing data from PSC lines, and Brian DeVeale for helpful discussions and critical review of the manuscript. We acknowledge the following funding sources: Helen Hay Whitney Foundation Postdoctoral Fellowship (RS), Ruth L. Kirschstein National Research Service Predoctoral Fellowship Award F31 HG011569-01A1 (TF), Weill Neurohub Fellowship (NKS), Fannie and John Hertz Fellowship (RAS), NSF Graduate Research Fellowship (RAS), National Institutes of Health DP2MH122400-01 (AAP), Schmidt Futures Foundation (AAP), Shurl and Kay Curci Foundation Innovative Genomics Institute Award (AAP), JSW is a Howard Hughes Medical Institute Investigator. AAP is a New York Stem Cell Foundation Robertson Investigator.

## Author contributions

R.S., T.F., J.S.W., and A.A.P. conceived the study design, executed all experiments, and wrote the manuscript. N.A.S. processed RNA-seq data and performed dN/dS calculations. R.A.S. provided the initial H1 CLYBL CRISPRi ES cell line and AAVS1 integration plasmids. B.J.P. provided high quality wildtype PSC lines and also provided extensive advice during the initial primary screens. J.S.W. and A.A.P. supervised all aspects of this work.

## Supplementary Figures

**Figure S1.**
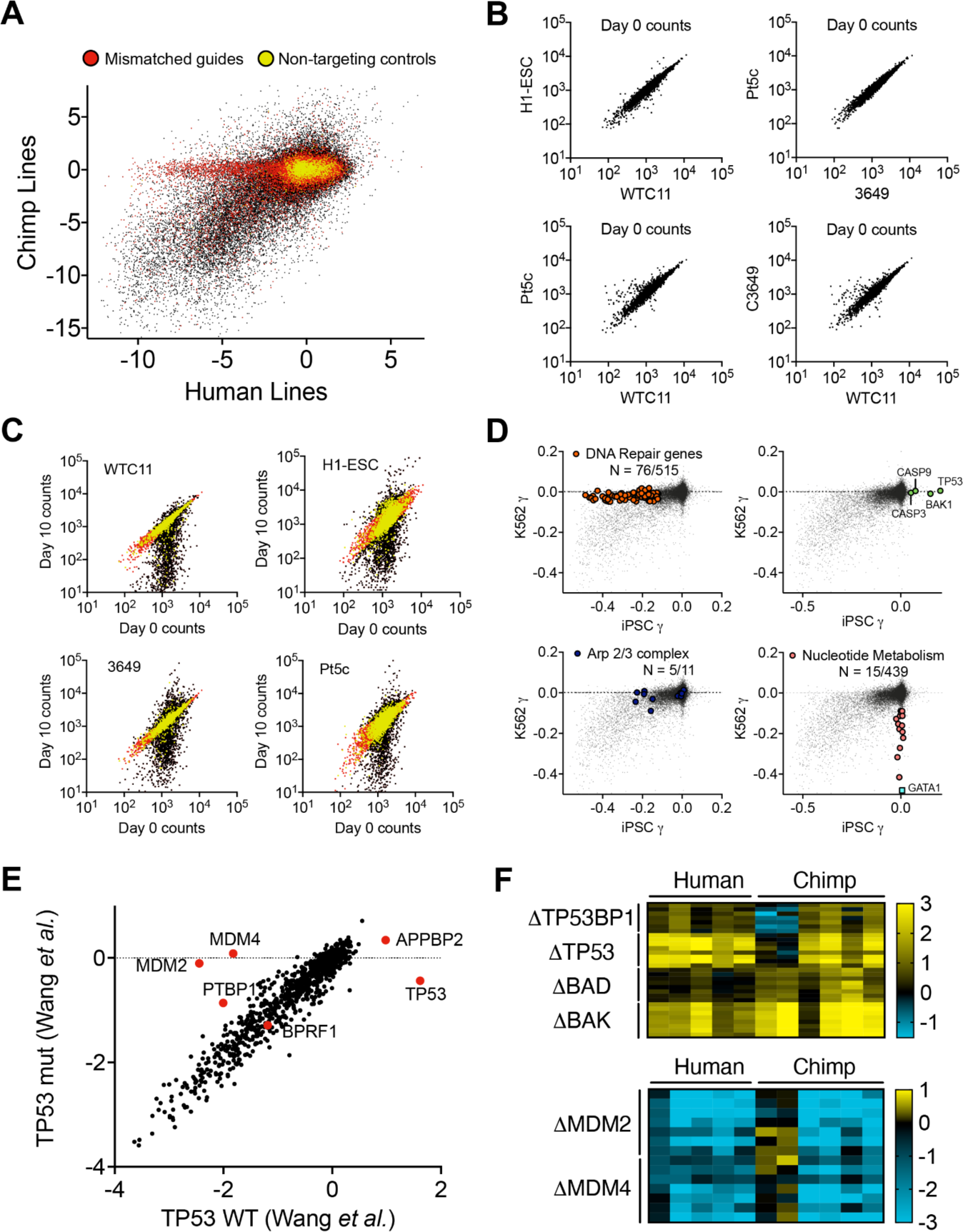
Genome-wide CRISPRi screens in human and chimpanzee PSCs, Related to Figure 1. (A) Log2 fold-change of sgRNA counts from genome-wide CRISPRi screens using the hCRISPRi-2 sgRNA library, averaged across two human and two chimpanzee cell lines. sgRNAs containing mismatches to the chimpanzee genome are colored in red and non-targeting sgRNAs are colored in yellow. Thus, a substantial number of mismatched sgRNAs targeting essential genes are depleted in human PSCs but not in chimpanzee PSCs. (B) Initial distribution of sgRNAs at growth day 0 (4 days post-infection). (C) Depletion or enrichment of sgRNA counts at growth day 10 compared to growth day 0. Non-targeting sgRNAs are colored in yellow, and sgRNAs characterized as non-significant by mean-variance modeling of a negative binomial distribution are colored in red. (D) Log2 fold-change of sgRNA counts averaged by gene (top 3) in PSC screens compared to prior screens in K562s^77^. (E) Log2 fold-change of sgRNAs averaged by gene from screens performed in TP53 wild-type and mutant acute myeloid leukemia (AML) lines^81^. Among the differentially essential genes targeted by the CEV-v1 sgRNA library, only ∼6 cause differential growth phenotypes in TP53 wild-type vs mutant cells (red) (F) Heatmap displaying log2 fold-change of sgRNA counts across five human and six chimpanzee PSCs, with columns 1, 6, and 7 showing primary genome-wide screening data and remaining columns showing data from secondary validation screening. Columns 6 and 7 (Pt5-C and C3649) represent the two chimpanzee PSCs that exhibit TP53 mutant phenotypes.

**Figure S2.**
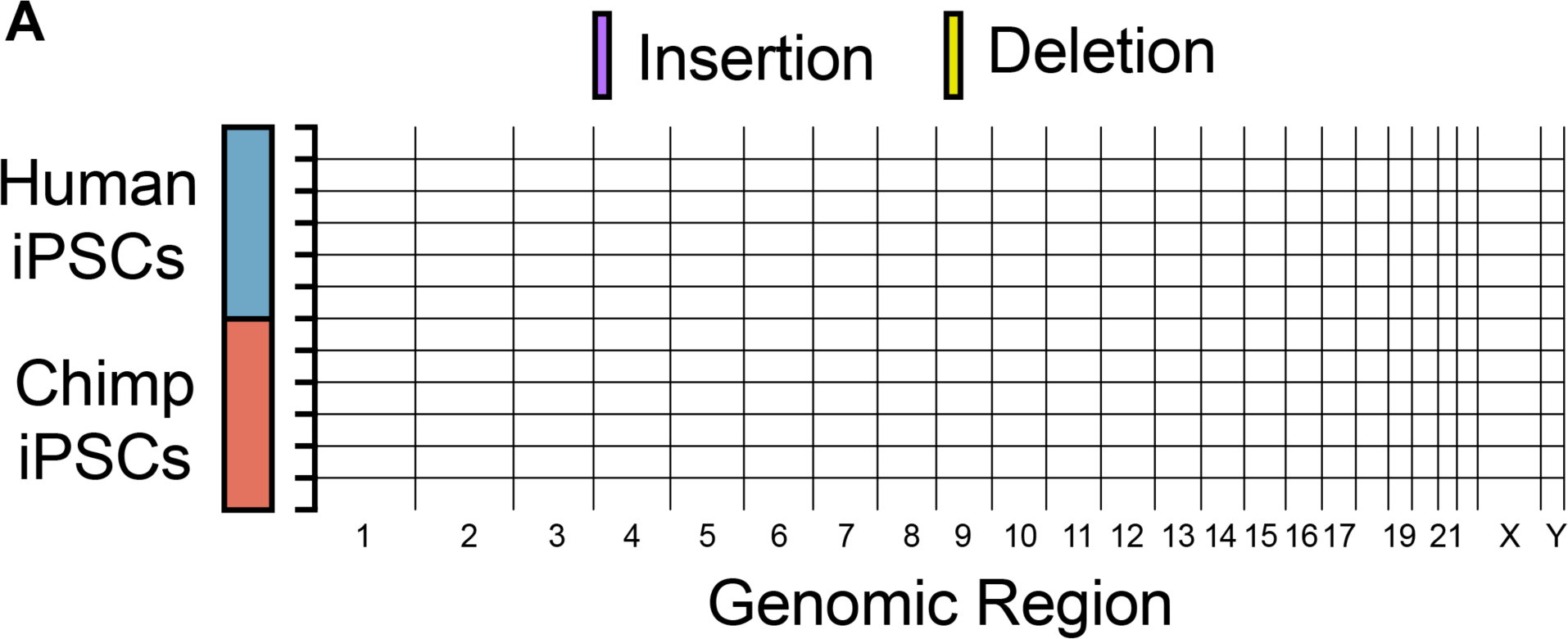
CEV-v1 validation screens in human and chimpanzee PSCs, Related to Figure 2. (A) CaSpER analysis of chromosomal copy number variations from bulk RNA-seq data across all newly engineered CRISPRi PSC lines as aligned to the human hg38 reference genome. For the chimpanzee genome, chromosome 2 refers to 2a and 2b.

**Figure S3.**
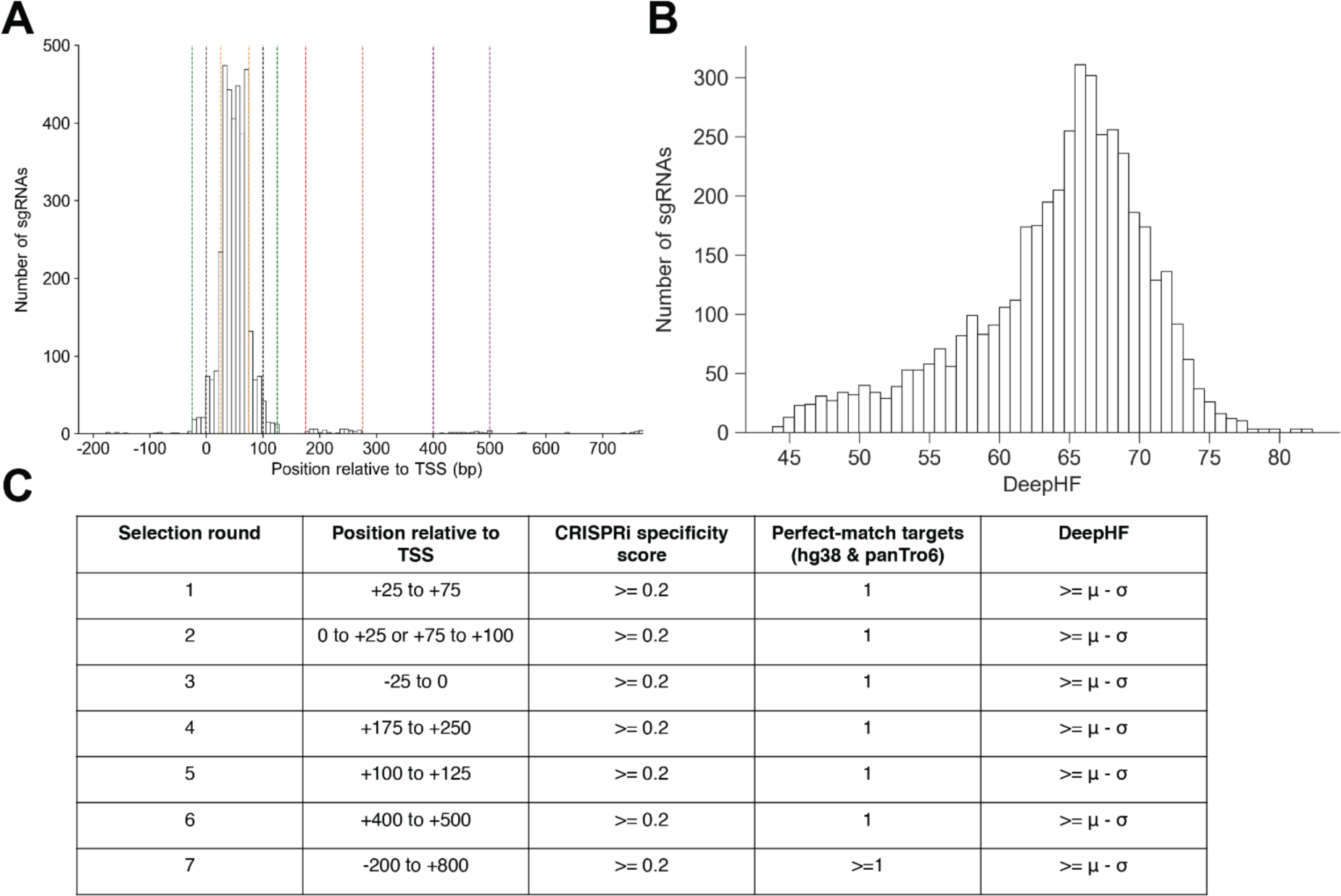
Design of CEV-v1 sgRNA library, Related to Figure 2. (**A**) Distribution of positions relative to the FANTOM-annotated TSS for sgRNAs in CEV-v1. Vertical colored lines indicate the selection round in which sgRNAs were chosen. (**B**) Distribution of DeepHF on-target predictions for sgRNAs in CEV-v1. (**C**) Selection criteria for CEV-v1 sgRNA library.

**Figure S4.**
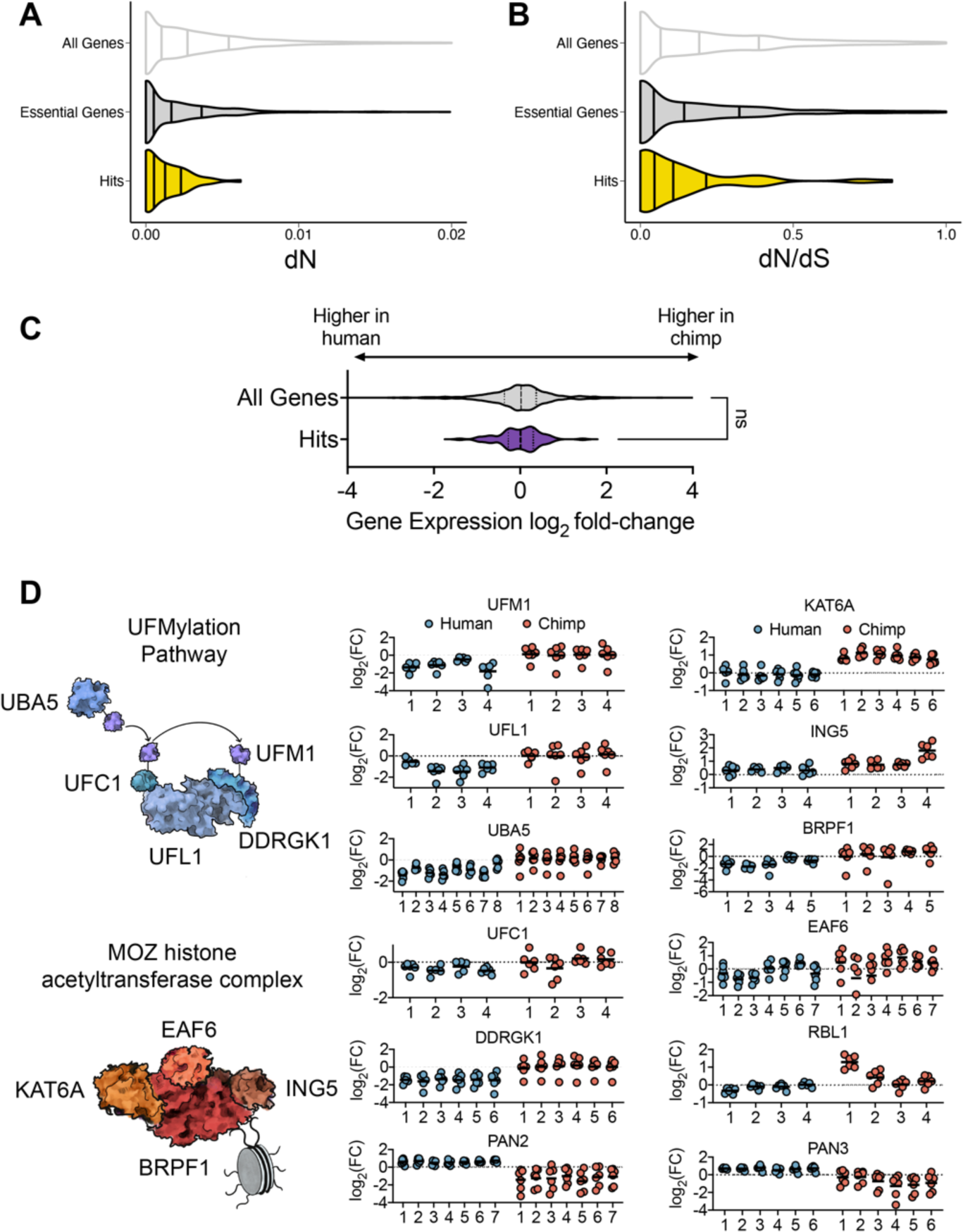
Species-specific genetic dependencies interact in biological processes and complexes, Related to Figure 3. (A) dN and dN/dS values for 75 validated differential-essentiality genes from this study compared to all genes or essential genes. (C) Comparative gene expression levels between human and chimpanzee PSCs for 75 validated differential-essentiality genes from this study vs. all genes expressed in PSCs. (D) sgRNA depletion or enrichment for all active sgRNAs targeting members of the UFMylation pathway, MOZ histone acetylation complex, RBL1, and the Pan2/3 complex. Each circle represents the sgRNA fold-change for one sgRNA in one human (blue) or chimpanzee (red) individual. Each stripplot contains a variable number of columns, corresponding to the number of significant sgRNAs targeting each gene. Genes with only one significant sgRNA (*ING5* and *RBL1*) are scored as less significant compared to genes with multiple significant sgRNAs and require validation of on-target effects.

**Figure S5.**
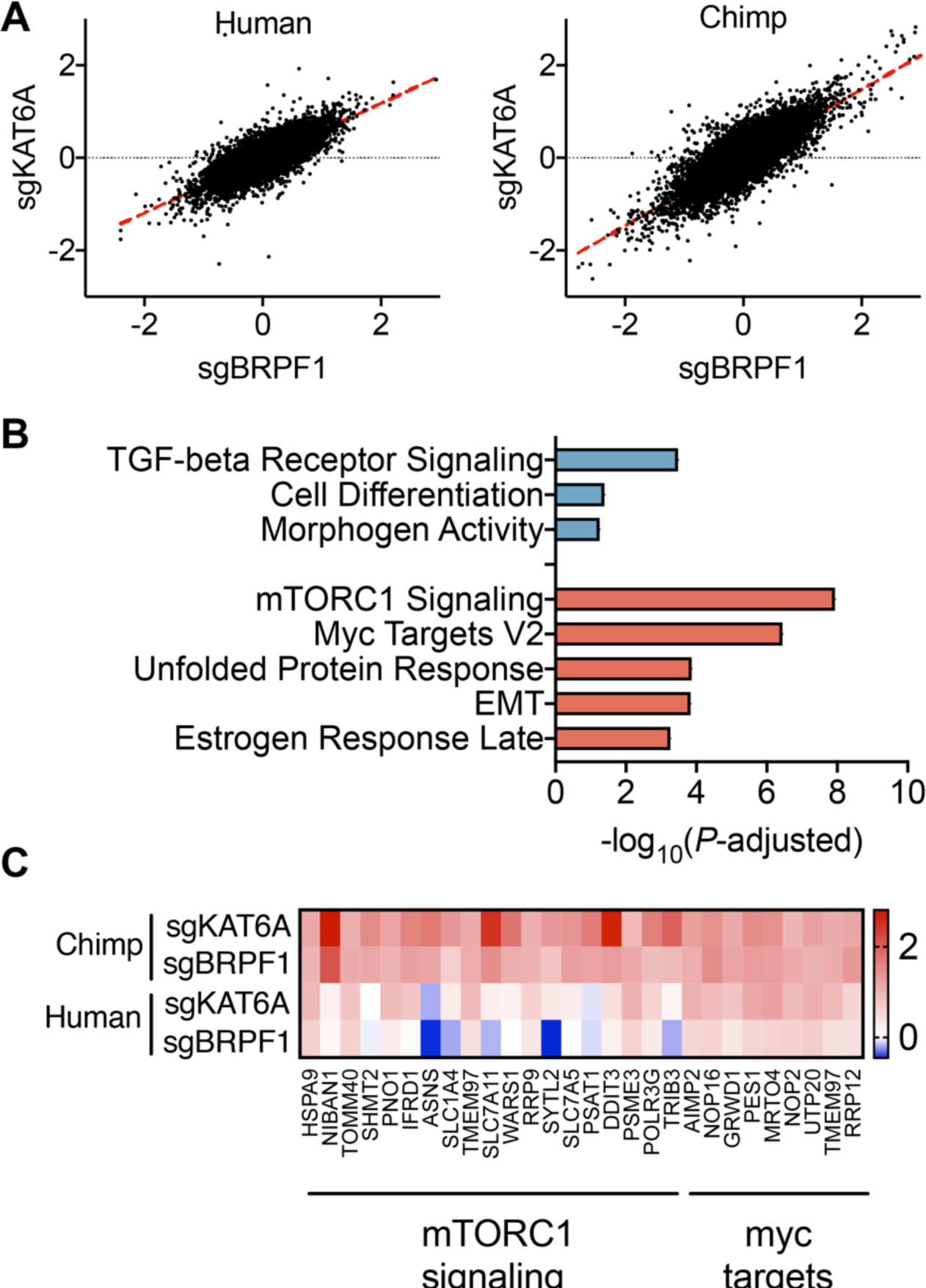
GO ontology enrichment for overexpressed genes in human and chimpanzee cells expressing sgKAT6A or sgBRPF1, Related to Figure 3. (A) −log10 adjusted *P*-values for overexpressed genes in human (blue) and chimpanzee (red) cells expressing sgBRPF1 or sgKAT6A. Overexpressed genes were defined as genes with fold-change > 2 in sgRNA expressing cells. Fold-change values were averaged between cells expressing sgBRPF1 and sgKAT6A. (B) Heatmap of RNA-seq expression data for human and chimpanzee cells depleted for KAT6A or BRPF1.

**Figure S6.**
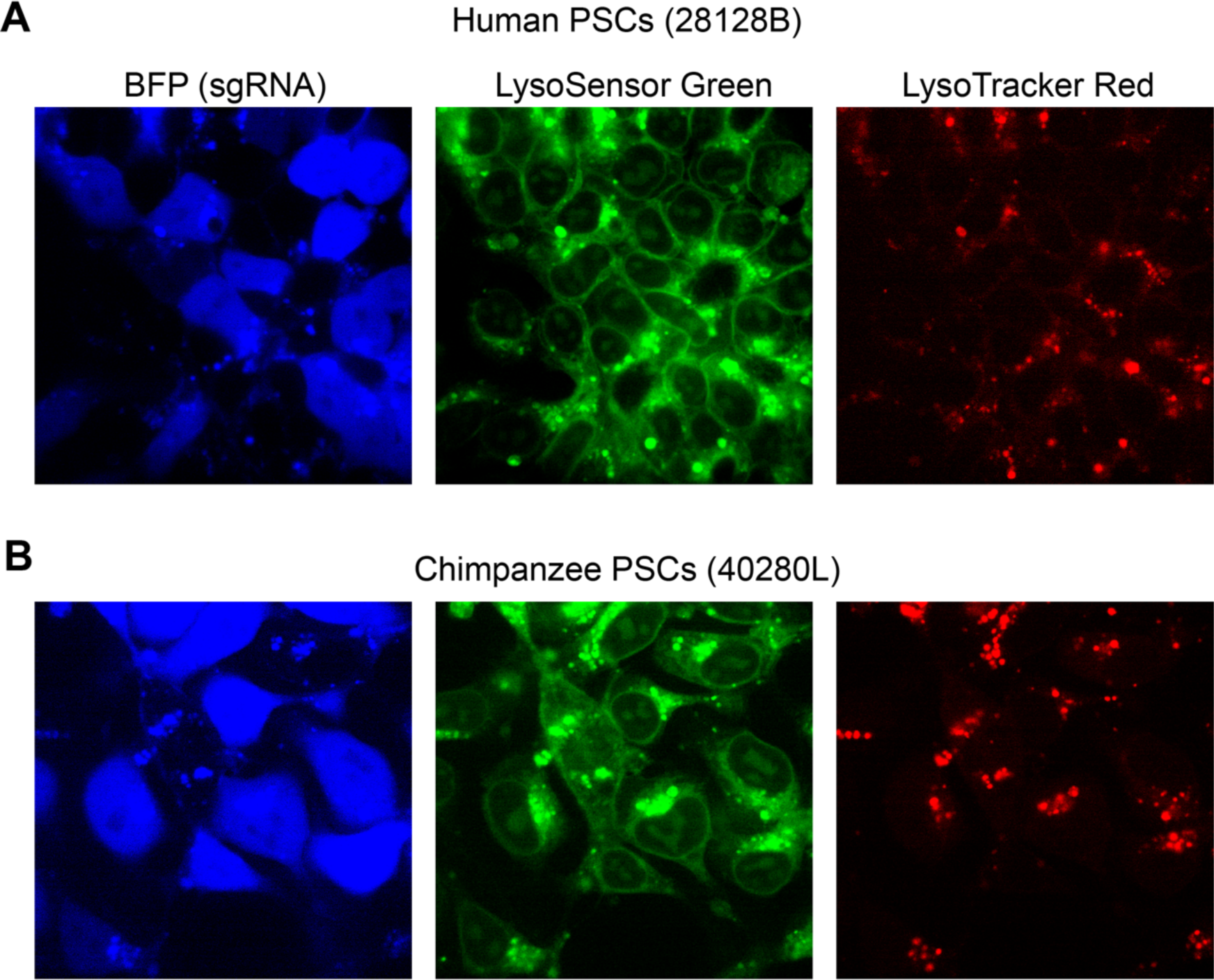
LysoTracker Red and LysoSensor green in *ATP6AP1* depleted cells, Related to Figure 3. (A) Co-culture of human PSCs (28128B) with wild-type cells (BFP-) and cells expressing sgATP6AP1 (BFP+) stained with LysoSensor Green and LysoTracker Red. (B) Co-culture of chimpanzee PSCs (40280L) with wild-type cells (BFP-) and cells expressing sgATP6AP1 (BFP+) stained with LysoSensor Green and LysoTracker Red.

**Figure S7.**
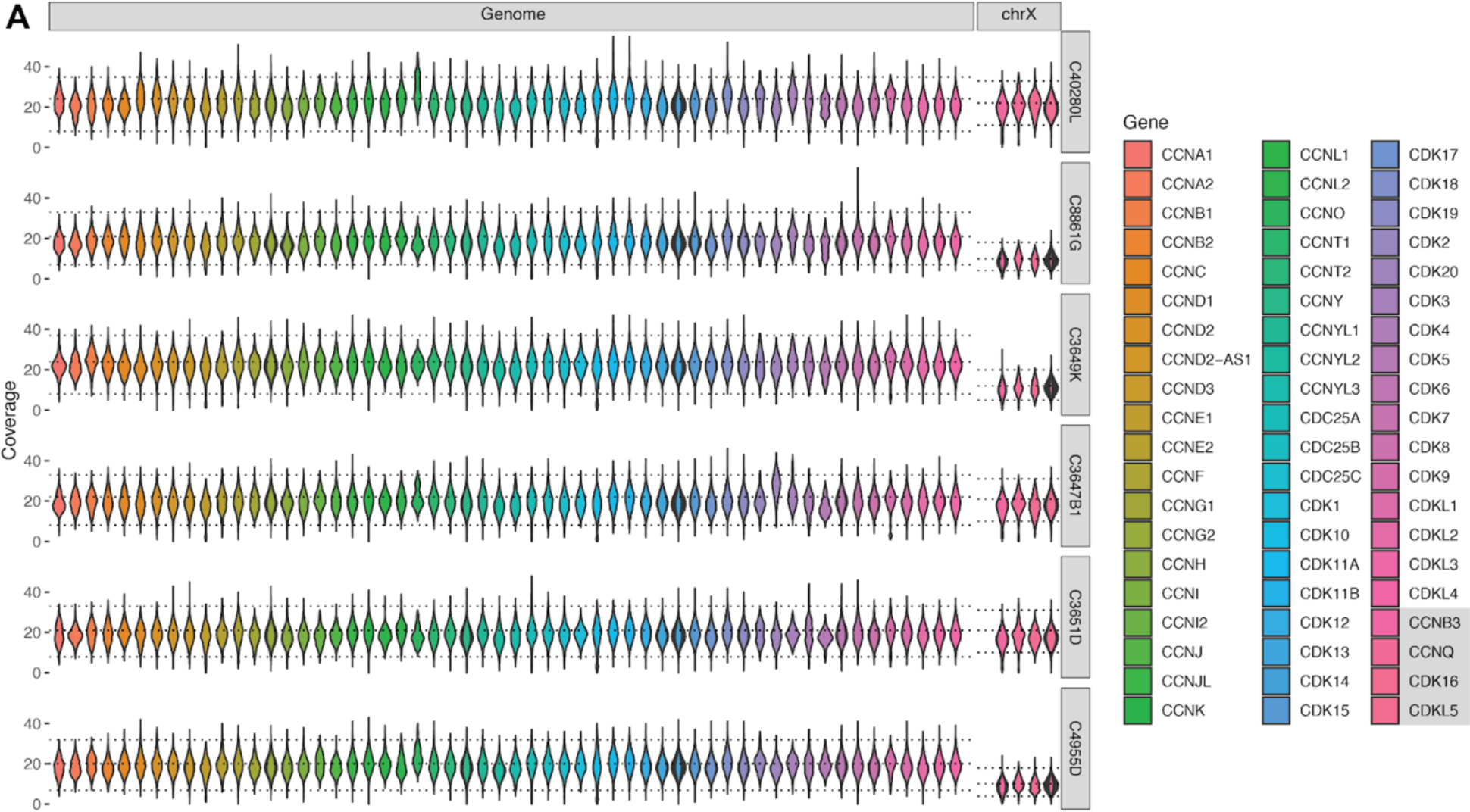
Analysis of whole-genome shotgun sequencing coverage at cyclin-CDK genes in chimpanzee PSCs, Related to Figure 4. (A) Whole-genome shotgun sequence coverage at all genes in the HUGO Gene Nomenclature Committee gene groups cyclins, cyclin dependent kinases, and class III Cys-based CDC25 phosphatases (https://www.genenames.org/). Each violin represents the coverage at each base across the entire body for each gene. The horizontal lines correspond to the 5th, 50th, and 95th percentiles of baseline coverage across the entire genome (“Genome” panel) or the X chromosome (“chrX” panel). Four genes in these sets located on chromosome X (CDKL5, CDK16, CCNB3, CCNQ) are shown separately to account for different baseline coverage; these gene names are outlined in a gray box in the legend. The top three rows correspond to chimpanzee PSCs from individuals used in the present study (C40280L, C8861G, C3649K), while the bottom three rows correspond to similarly reprogrammed chimpanzee individuals.

**Figure S8.**
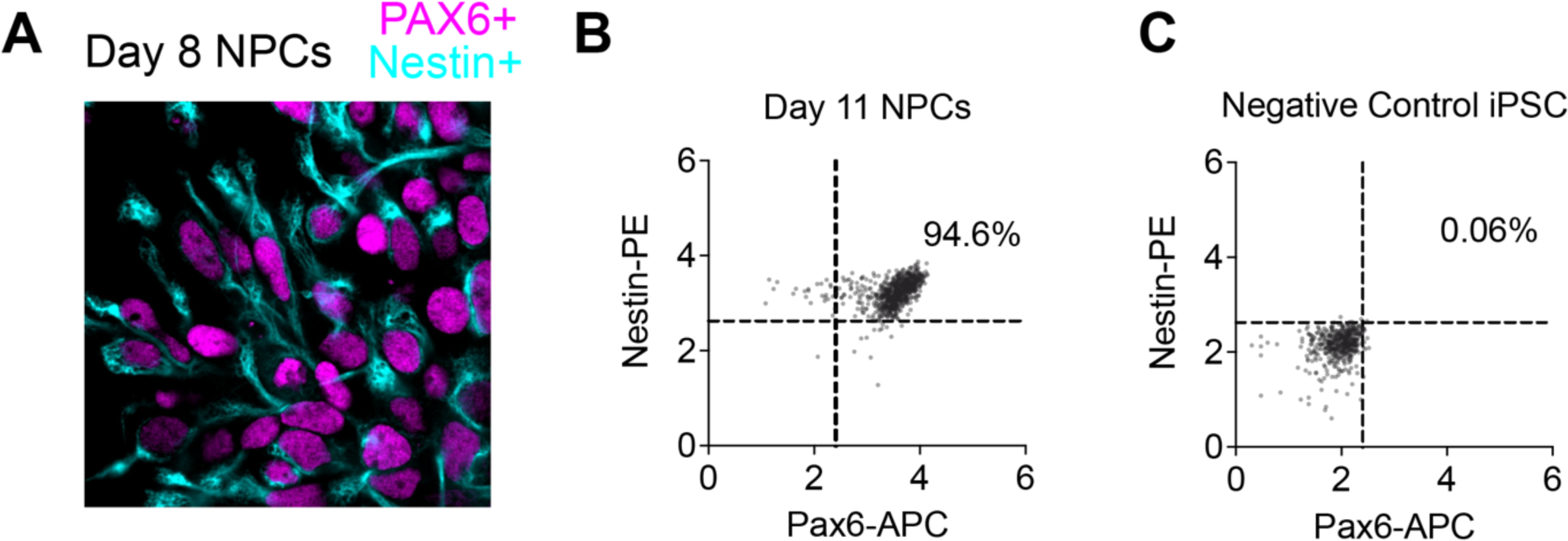
Derivation of human and chimpanzee NPCs, Related to Figure 5. (A) Chimpanzee neural progenitor cells (C40280L) stained for Pax6 and Nestin, visualized by confocal microscopy. (B) Chimpanzee neural progenitor cells (C40280L) stained for Pax6 and Nestin, quantified by flow cytometry. (C) Negative control PSCs stained for Pax6 and Nestin, quantified by flow cytometry.

**Figure S9.**
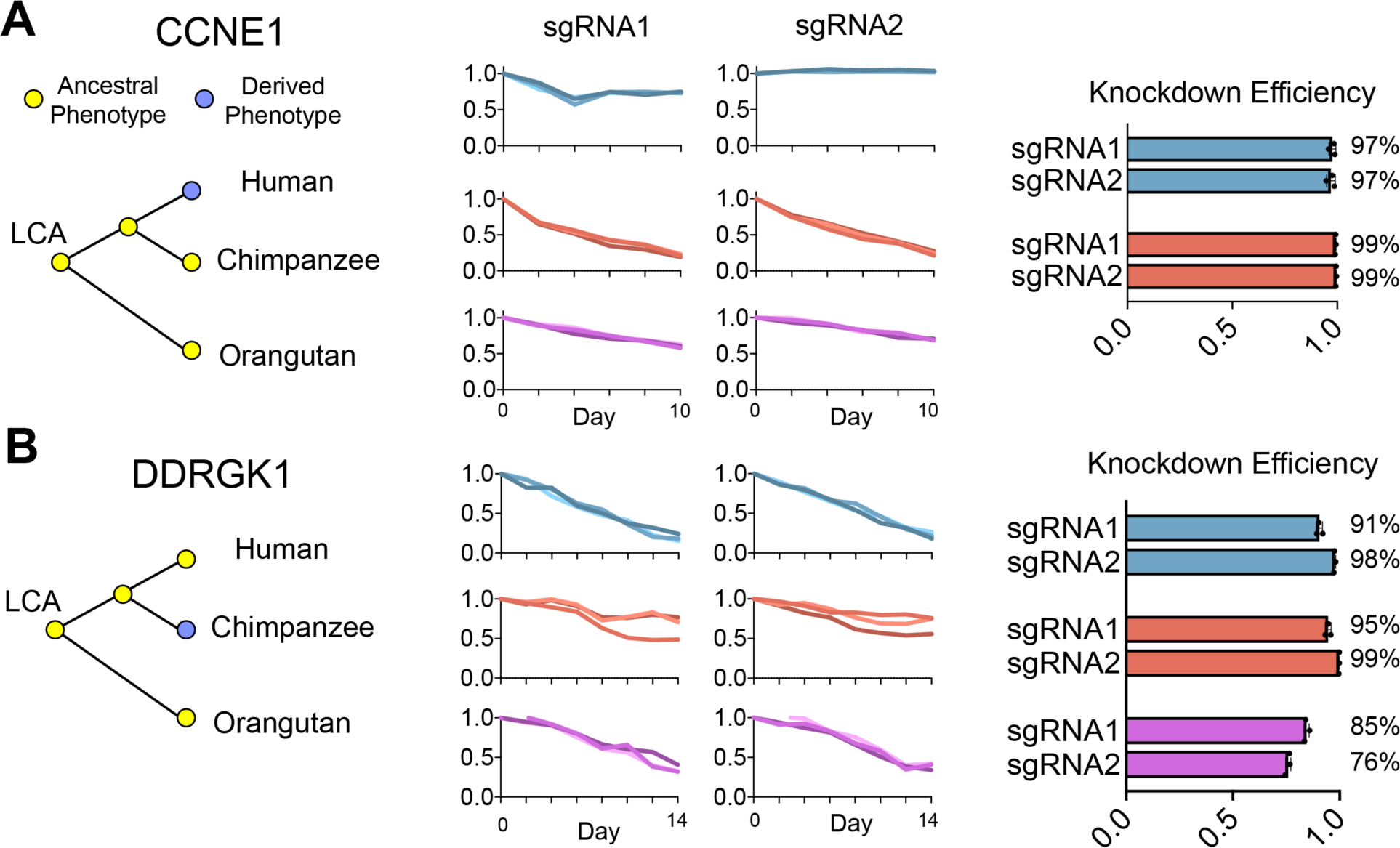
Tri-species comparison of human, chimpanzee, and orangutan PSCs expressing sgRNAs targeting *CCNE1* and *DDRGK1*, Related to Figure 6. (A-B) Change in the relative fraction of CCNE1 (A) and DDRGK1 (B) sgRNA containing cells over time in human, chimpanzee, and orangutan PSCs. qRT-PCR measurements of sgRNA knockdown efficiency for each sgRNA.

**Table S1. Cellular metadata for primary CRISPRi screen and CEV-v1 screens**

**Table S3. sgRNA counts from hCRISPRi-v2 genome-wide screens**

**Table S4. hCRISPRi-v2 sgRNA library**

**Table S5. sgRNA counts from CEV-v1 validation screens**

**Table S6. qRT-PCR primers and individual sgRNA primers**

**Table S7. Transcriptional responses to BRPF1 and KAT6A repression in human and chimpanzee PSCs**

